# scCont: Interpretable Contrastive Learning for Functional Characterization of Cellular State Transitions

**DOI:** 10.64898/2026.11.05.764599

**Authors:** Neal Kewalramani, Indranil Paul, Andrew Emili, Stefan Wuchty, Mark Crovella

## Abstract

Timecourse single-cell RNA (scRNA) datasets have become indispensable biological tools for studying and uncovering the underlying regulatory processes driving continuous cell-state transitions such as the epithelial-to-mesenchymal transition. However, these highly fluid, non-linear gene program dynamics remain difficult to disentangle. While deep neural network models can effectively map complex topologies, they are biological black boxes, lacking the gene-level interpretability required by experimental practitioners. Here, we introduce scCont, a fully unsupervised contrastive learning framework for extracting interpretable functional transition units from scRNA-seq data. Rather than relying on explicit temporal labels or clustering priors, scCont uses an unsupervised k-nearest neighbors approach to project local transcriptomic neighborhoods into a low-dimensional embedding. During post-training analysis, scCont employs Shapley network attribution to determine gene-latent relationships, mapping latent dimensions directly into functional gene programs. We quantitatively and qualitatively validate this representation by demonstrating that scCont captures biologically interpretable dynamics more effectively than baseline methods such as LDVAE. Evaluated across 13 diverse EMT timecourse datasets and four simulated datasets, scCont exposes interpretable functional dynamics and effectively maps context-dependent functional groups, helping to bridge the gap between deep learning representation and translational biological utility. scCont is an open source software available on GitHub (https://github.com/nramani611/scCont).

## Introduction

Timecourse scRNA datasets are essential tools for capturing dynamic biological processes across timepoints. The gene programs encapsulated in these datasets reveal how cells evolve in response to stimuli, during the cell cycle, or throughout disease progression and cellular reprogramming. However, cell state transitions are often complex: individual cells often exhibit heterogeneous developmental rates and branching of cell fates frequently occurs, complicating the disentanglement of dynamic gene modules.

In particular, the epithelial-mesenchymal transition (EMT) is a prime example where asynchronous transitions and intermediate states challenge traditional linear analysis methods. Specifically, EMT is a key biological process involved in many cancers, during which cells lose their stationary, epithelial identity to adopt a more motile, mesenchymal cell state, promoting cell migration, cancer invasion, and tumorigenesis. This transformation is characterized by the rewiring of many subcellular and extracellular processes, such as the actin cytoskeleton, the extracellular matrix, and cell survival pathways (1). These gene programs, however, are difficult to characterize in the intermediate state (E/M), where cells exist in a hybrid, transient phase between the epithelial and mesenchymal states (2). Furthermore, EMT is highly context-dependent; different cell lines and inducers activate different gene networks (3). To systematically study such a complex system, EMT needs to be considered as a collection of context-adaptive gene programs that can dynamically reconfigure rather than a single continuum.

To investigate the E/M state, scRNA-seq timecourse experiments are frequently employed to capture transcriptional dynamics. With these high-dimensional timecourse datasets, dimensionality reduction remains a primary method for distinguishing biological signals from experimental noise. While linear models such as principal component analysis (PCA) and non-negative matrix factorization (NMF) (4) are frequently employed for extracting linear gene programs, and form the core of current analytical tools (5; 6), such methods often struggle to map the highly non-linear, temporally continuous program dynamics arising during cellular state transitions.

Recently, neural networks have gained traction in scRNA-seq analysis as a result of their ability to capture non-linear gene interactions (7). However, neural architectures generally lack interpretability of their latent features, preventing the extraction of biological mechanisms (8) that are more readily achievable with linear methods like PCA and NMF. While neural networks are widely used in broader omics applications (9), improving their interpretability would amplify their utility for scRNA-seq analysis.

To bridge this gap, we turn to the framework of contrastive learning (10; 11). While contrastive learning is a natural tool for denoising and dimensionality reduction, its use in scRNA analysis has generally followed the *data augmentation* paradigm (12; 13), a strategy in which the positive pairs used for learning are constructed by perturbing data. Thus, it has not in general been used to generate interpretable outputs for timecourse data.

Here, we achieve interpretable and temporal dimensionality reduction for timecourse scRNA data via scCont, an unsupervised contrastive learning framework designed to map cell-state transitions. The core innovation of scCont is its unsupervised, geometry-aware architecture, combined with its use of attribution methods to achieve interpretability. Its geometry-based approach uses a *k* - nearest neighbor (*k* -NN) graph to dynamically identify positive pairs based strictly on local temporal neighborhoods. This approach projects high-dimensional neighborhood relationships into a low- dimensional embedding, preserving biological continuity without relying on explicit temporal labels or linear assumptions. We couple that framework with a rigorous attribution stack to translate latent features into highly interpretable, statistically defined gene modules. We demonstrate the utility of scCont across a variety of timecourse datasets for EMT under different conditions. We first validate scCont on four SERGIO (14) simulated datasets, confirming its accuracy based on known ground truth. We then demonstrate the effectiveness of scCont by applying it to an MCF10A (breast epithelial) cell line treated with TGFB1 (15). Finally, we illustrate scCont’s ability to resolve context-dependent gene programs and cross-dataset module alignment by analyzing 12 additional EMT datasets from Cook et al. (3). Across these contexts, we demonstrate that scCont uncovers temporally evolving processes that elucidate how experimental context, inducer type, and genetic background dynamically reconfigure the E/M state.

## Related Work

Recent advancements in single-cell transcriptomic analysis can be broadly categorized by their approach to balancing model complexity with biological interpretability.

### Linear Representations and Matrix Factorization

Historically, linear dimensionality reduction methods have been widely utilized to analyze scRNA-seq datasets. NMF, in particular, extracts factorized matrices that provide a clear, additive connection to underlying biological programs (16). To enhance stability and local topology preservation, several advancements have been introduced. For example, consensus NMF (cNMF) runs NMF over many initializations and clusters the resulting factors together to robustly identify gene sets (5), while Laplacian regularization methods have been developed to improve NMF performance by preserving local geometric structures within scRNA data (17). However, despite these improvements and their high interpretability, NMF architectures remain fundamentally optimized to learn parts-based representations. As demonstrated by models such as scRCMF and foundational work by Shao & Höfer (18), NMF excels in identifying distinct cell type populations, emphasizing its parts-based nature, but struggles to quantitatively characterize cells in highly plastic transition states (19). Consequently, these linear methods often force fluid biological processes—such as continuous, developmental functional groups—into discrete, static subpopulations. We illustrate this effect in our results.

### Deep Generative and Contrastive Models

Deep learning frameworks have emerged as powerful tools for modeling complex, nonlinear transcriptomic distributions. For example, variational autoencoders (VAEs), such as scVI (20), have established the standard for modeling the complex, nonlinear distributions in underlying scRNA-seq data. Similarly, generative models like scGen successfully predict out-of-sample single-cell perturbation responses (21); however, the authors themselves note that extracting the specific biological mechanisms driving these divergent responses remains a computational challenge (8). To address this “black box” limitation, interpretable factor models such as linearly decoded variational autoencoder (LDVAE) (22) were developed to map gene weights directly to latent features. Yet, this interpretability constrains the decoder to linear mappings, sacrificing significant nonlinear representational accuracy. In contrast, scCont leverages authentic spatial and temporal neighborhoods—rather than artificial augmentations or discrete priors—to map continuous, nonlinear gene program fluxes, bridging the gap between topological mapping and biological interpretability.

### Contrastive Learning for scRNA

Contrastive learning has seen many applications in the scRNA-seq data analysis field. Established methods, such as CLEAR, Concerto, and scContrast (12; 13; 23), typically use standard data augmentations—like random gene dropout and Gaussian noise—to extract meaningful latent features from gene expression vectors, but they are not optimized for mapping dynamic state transitions. More recently, scCRT (24) applies contrastive learning to trajectory inference, yet it relies on prior clustering information to generate positive pairs, effectively reinforcing discrete state boundaries. While other contrastive models like scGCL employ a nearest-neighbor selection method for data imputation (25), none of these existing approaches attempt to bridge the interpretability gap or map continuous cell state transitions in timecourse scRNA-seq data.

## Results

### scCont Algorithm Overview

Unlike trajectory inference methods, scCont does not order cells along a pseudotime axis; instead, it learns latent features that capture coordinated shifts in functional gene programs across conditions or timepoints. Figure 1 shows an overview of scCont applied to scRNA- seq data. The key output of scCont is a set of low-dimensional, temporally-sensitive latent features that are interpretable in terms of biological functions. We refer to these latent features as functional transition units (FTUs) to reflect the biological and temporal information they encapsulate. We then cluster these latent features based on their spatial variation to identify modular transition units. These modular transition units describe how the latent feature clusters break up the biological pathways into distinct, interpretable processes.

**Figure 1.**
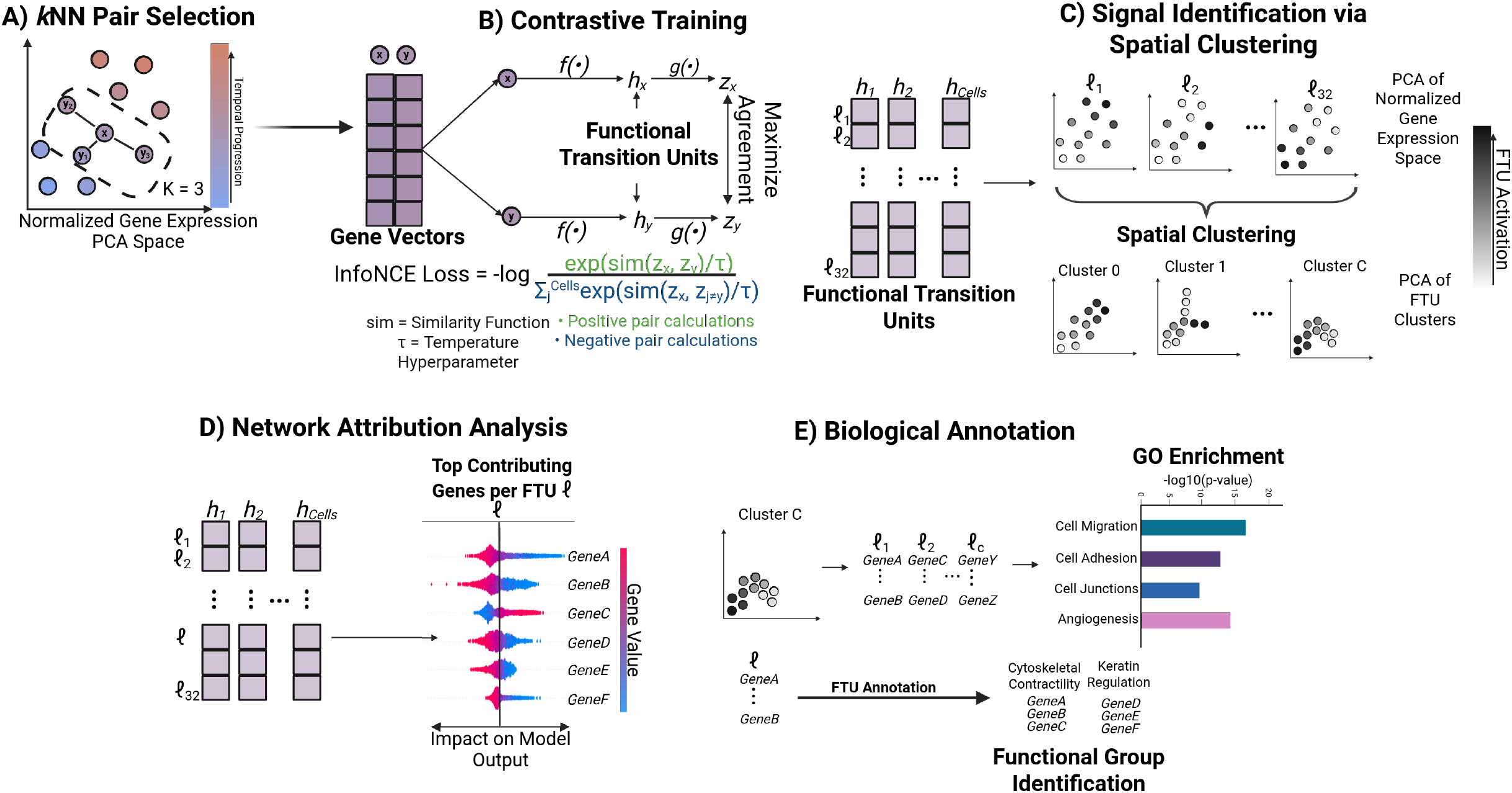
Overview of the scCont algorithm. The framework operates in five stages to extract modular functional transition units (FTUs) from timecourse scRNA-seq data. FTUs are defined as a latent unit that captures the coordinated shift in functional gene programs across different conditions or timepoints. **A) k-NN Pair Selection:** A *k*-Nearest Neighbor graph (k=3) is applied to the normalized gene expression space to dynamically identify positive pairs. PCA is used here to reduce noise in the data. **B) Contrastive Training:** These positive pairs serve as inputs to a contrastive learning architecture that minimizes InfoNCE loss to maximize agreement between topologically adjacent cells in the latent space. *f(•)* refers to the encoder that generates the FTUs (*h*), and *g(•)* refers to the projector function that makes the similarity embeddings (*z*). Figure adapted from Chen et al (10). **C) Signal Identification via Spatial Clustering:** FTU activations are projected onto the gene expression PCA space; features with correlated spatial patterns are grouped to isolate distinct signal modules. **D) Network Attribution Analysis:** Specific latent-defining genes for each FTU are identified using feature importance scores (SHAP). **E) Biological Annotations:** FTUs are connected to biological processes via a multi-tiered validation approach, combining broad Gene Ontology (GO) enrichment for clusters with functional group identification for individual top-contributing genes.

Intuitively, the contrastive learning framework seeks to create representations that drive the representations of similar pairs of cells together and dissimilar pairs apart in latent space (Fig. 1B). The model, therefore, is forced to learn relationships between data points (and not just learn representations of data points independently, as do autoencoders). Contrastive learning operates by training two models simultaneously: an encoder function (*f(•)*) and a projector (*g(•)*). The goal of the encoder function is to create the low-dimensional embedding of the input data; these latent embeddings are used by the projector to train the contrastive learning model using InfoNCE loss (11). The loss calculations shown in Figure 1B show how the InfoNCE loss drives contrastive learning. The result of this training method is an encoder function that projects the scRNA data into embeddings focusing on the subtle differences between samples, making it a suitable tool for identifying FTUs. A key novelty in scCont is in defining the positive pairs: scCont is specifically designed in an unsupervised manner to reduce dependence on additional label data and preserve local cell-cell relationships. Figure 1A illustrates the use of a *k* -NN model to identify positive pairs. Here, scCont uses the local neighborhood information supplied by *k* -NN to train and encode the high-dimensional temporal neighborhoods into an FTU.

To illustrate the effectiveness of *k* -NN positive pair selection for extracting temporal information, we perform an ablation study (Supplementary Fig. 4A) with the MCF10A TGFB1 dataset. We find that standard data augmentation methods for contrastive learning fail to capture most timepoint information. As a result, derived latent features do not encode information necessary to track functional gene programs. However, under *k* -NN sampling of positive pairs, latent features are effective at encoding temporal evolution. We also assess how *k* -NN incorporates temporal evolution by measuring the number of within-timepoint pairs as a function of *k* (Supplementary Figure 5C).

To identify modular transition units (MTUs) from scCont, we perform spatial clustering of the FTUs (Fig. 1C). This specific step allows us to separate latent features representing different spatial patterns in the scRNA data. We define an MTU as a group of FTUs sharing similar temporal dynamics and spatial co-localization in PCA space. The purpose of this step is to characterize the different pathways taken by the genes that are represented as latent features. We provide an example of this in Figure 3. Clustering is performed by projecting FTUs into this normalized gene expression PCA space and identifying where they co-localize, revealing groups with coherent spatial patterns and consistent biological annotation via Gene Ontology (GO) enrichment (see Methods for more details). While MTUs provide broad, biological trends, we analyze genes on an FTU level to identify specific functional groups and their changes in scRNA data. To interpret these learned FTUs, scCont employs SHapely Additive exPlanations (SHAP), which maps the non-linear relationships between gene expression levels and output latent space. For each gene, SHAP calculates a marginal contribution score, evaluating how the FTU changes when the gene’s contribution is withheld across all possible gene combinations. SHAP enables specific genes to be attributed to driving local, latent state transition behavior, making the algorithm a suitable interpretability method for identifying FTUs in timecourse scRNA-seq data (26; 27). We isolate the most significant contributing genes for each FTU using a defined attribution threshold (*z >* 2.57, *p <* 0.01) based on mean absolute SHAP values. These statistically derived gene sets are tested for Gene Ontology (GO) enrichment (*adj. p-val <* 0.05) to map the FTUs to high-level biological processes. FTUs are then characterized by the highest scoring genes within each set of significant GO terms. We then contextualize these SHAP derived genes in the literature (Fig. 1E) to validate their known regulatory roles (full gene annotations and literature search results are available in Supplementary Materials). This multi-tiered approach ensures that scCont outputs are robust, reproducible, and biologically interpretable. To ensure that the FTUs extracted from scCont reproduce similar, significant gene sets we conduct a SHAP seed stability test. By measuring the Jaccard value of gene sets across five random initializations on the SHAP step, we found that gene sets from most FTUs are highly stable (Supplementary Figure 4B).

To validate scCont’s ability to capture and align FTUs across varying contexts, we apply it to a variety of simulated datasets as well as EMT timecourse scRNA-seq datasets. First, we demonstrate scCont’s capacity to recover ground-truth functional groups by applying the model to four SERGIO (14) simulated datasets. Next, we showcase scCont’s capability to disentangle the EMT continuum in an MCF10A cell line (breast epithelial) treated with TGFB1. This data was collected over the course of 8 days during which cells were sequenced daily (Days 0-7) (15). In addition, to demonstrate the framework’s ability to model cross-dataset module alignment and behavior across different stimuli, we analyze 12 datasets from Cook et al. (3) encompassing four cell lines: A549 (lung), DU145 (prostate), MCF7 (breast), and OVCA420 (ovarian). Each cell line was treated with three distinct inducers: tumor necrosis factor (TNF), TGFB1, and epidermal growth factor (EGF). The experiments ran for 8 days during which cells were sequenced at 0h, 8h, 1d, 3d, and 7d, followed by inducer withdrawal to capture state reversion (8h off, 1d off, 3d off). Analyzing this diverse array of conditions demonstrates how specific FTUs adaptively reconfigure depending on the genetic context and inductive signal, establishing the framework’s broader utility in resolving context-dependent state transitions.

### scCont disentangles ground truth functional programs from noise in simulated data

In this section, we conduct a sensitivity analysis to analyze the performance of scCont as a function of the hyperparameter *k*. Furthermore, we compare the performance of scCont against other baseline models to see how the model performs against the state- of-the-art. Using SERGIO, we created four simulated datasets with the following characteristics: a bifurcation (two lineages; 800 program genes and 16 functional programs with 50 program genes per functional group; 2200 background genes), a trifurcation (three lineages; 800 program genes and 16 functional programs with 50 program genes per functional group; 2200 background genes), at maximum latent capacity (two lineages; 800 program genes and 32 functional programs with 25 program genes per functional group; 2200 background genes), and a diluted signal (two lineages; 400 program genes and 16 functional programs with 25 program genes per functional group; 2600 background genes). SERGIO is a scRNA data generator that simulates gene expression with input gene regulatory networks. We designed the input networks to SERGIO that simulate the expression of our input functional groups (see Methods for details). The background genes are of the same distribution of the program genes but randomly permuted across the cells; this process ensures that the background genes are viable distractions but lack any program structure. We validate scCont across all four of these different signals to illustrate that the model can disentangle the functional groups across these different simulated conditions. We highlight the bifurcation dataset in the main text, where the established functional groups delineate into two branches in Figure 2 and the other simulated datasets in Supplementary Figures 1 and 2. Figures 2A-D illustrate that scCont resolves known lineage structure. Projecting all 3000 genes onto the PCA space leaves the two lineages intermixed (Fig. 2A). We demonstrate the ground truth structure by only performing PCA on the known 800 program genes in Figure 2B. It is important to note that the program genes would not be known in practice but serves as a comparison with Figure 2C to illustrate that scCont resolves a similar structure. Trained with neighbors from Figure 2A, the scCont model resolves a structure similar to the ground truth structure in Figure 2B (Fig. 2C). We show these visualizations to emphasize the fact that scCont is capable of extracting the important program genes from a noisy dataset, while also resolving the ground truth lineage of the cells.

**Figure 2.**
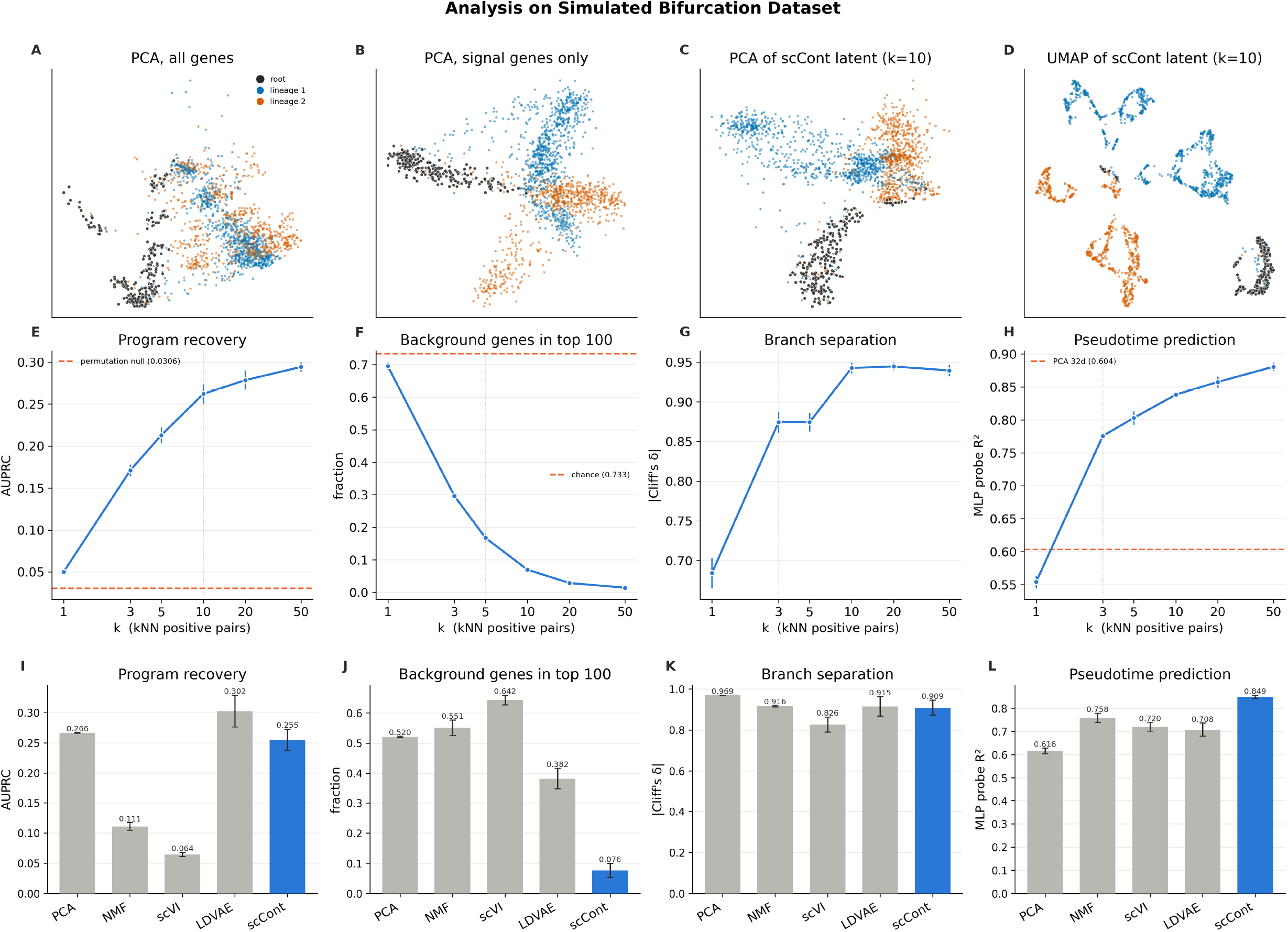
scCont analysis on bifurcating simulated dataset. Simulated bifurcating dataset generated with SERGIO with 800 program genes assigned to 16 functional programs and 2200 background genes. **Panels A-D)** The same cells are presented in four different spaces and are colored by ground truth lineage (lineage refers to the different branches in the bifurcation). **A)** Principal component analysis (PCA) of all 3000 genes. **B)** PCA of the 800 program genes only. **C)** PCA of the 32-dimensional scCont latent space (*k* = 10). **D)** UMAP of the scCont latent space. **E-H)** Sensitivity analysis of scCont as a function of *k*-Nearest Neighbors (ran over five random seeds). **E)** Hungarian-matched AUPRC of program genes under SHAP ranking of each program’s best-matched latent. **F)** Fraction of each latent’s top 100 highest-attributed genes belonging to no program. Top 100 genes are considered to ensure a consistent comparison across random seeds. **G)** Largest absolute Cliff’s *δ* between lineages across latent dimensions. **H)** Cross-validated *R*^2^ of an MLP probe predicting ground truth pseudotime from embedding. **I-L)** Comparison of scCont (*k* = 10) against state-of-the-art methods (all models are at 32 latent features). **I)** AUPRC of program recovery. Hungarian matching is implemented to find the best one-to-one pairing of one program and one latent feature. **J)** Average proportion of background genes in top 100 contributing genes for each latent feature. Top 100 genes are considered to ensure a consistent comparison across all models. **K)** Cliff’s delta (*δ*) between lineages to measure how well each model distinguishes between ground truth lineages. **L)** A multilayer perceptron (MLP) probe is used to predict pseudotime (*R*^2^) from the latent features from each model as a measure of temporal preservation.

Figures 2E-H test four different metrics as a function of *k*. At *k* = 10, matched program recovery reaches an AUPRC of 0.26; background genes make up 7.0% of the 100 highest-attributed genes per latent, a ten-fold depletion relative to the 73.3% chance level; sister lineages separate with a maximum absolute Cliff’s *δ* of 0.94; and an MLP probe recovers ground-truth pseudotime from the embedding at *R*^2^ = 0.84, against 0.60 for a 32-component PCA of the same matrix. The saturation point varies modestly across the other three simulations in Supplementary Figure 1. Based on these results, we conclude that scCont’s architecture is capable of disentangling the program signal. Specifically, we observe program recovery, the reduction of background noise, and the separation of lineages that are beyond chance.

To put scCont’s performance in proper context, we evaluate the performance of four other models and compare scCont against these other methods. For each model, we measure the four statistics in Figures 2E-H for other baseline state-of-the-art methods. Figure 2I- L establishes scCont as a top performer in each measured metric when compared against the four other baseline methods. PCA is the canonical linear baseline, and NMF forms the basis of many interpretable scRNA-seq models (28; 29). LDVAE pairs a non-linear encoder with a linear decoder specifically to recover interpretable gene loadings (22). scVI serves as a state-of-the-art baseline for the latent representation of scRNA-seq data (20). For scVI, we add the SHAP attribution analysis post-training, similar to scCont, to differentiate the results we see as evidence of the contrastive framework instead of purely the SHAP attribution analysis. In terms of program recovery, LDVAE slightly outperforms scCont for this simulated dataset but the two remain comparable (Fig. 2I), but we find that this result can change based on the simulation conditions. In Supplementary Figure 2A, E, and I, we demonstrate how this result slightly changes based on simulation conditions; ultimately, depending on the conditions, scCont is either comparable or better at program recovery compared to LDVAE. What remains consistent, however, is the fact that scCont is much more resilient to background noise compared to other models (Fig. 2J and Supplementary Fig. 2B, F, J) and maintains a signal that represents the ground truth temporal structure in the data (Fig. 2L and Supplementary Fig. 2D, H, L). We attribute this result to scCont’s *k* NN pair selection method (see the ablation study in Supplementary Figure 4A for additional results). By creating an embedding through the comparisons of nearby data points, scCont effectively resolves the program genes and the temporal structure of the functional programs in a completely unsupervised manner. All models perform well in terms of distinguishing the two known branches (Fig. 2K and Supplementary Fig. C, G, K). Overall, we conclude that scCont is a top performer in program recovery and is the best performer in noise reduction and temporal recovery compared to baseline methods.

The analysis on simulated data supports our claim that scCont’s latent features can be interpreted as functional transition units (FTUs). Specifically, we show that the algorithm is capable of identifying genes from the same functional group, while excluding those that belong to none. We also demonstrate that the same FTUs carrying those programs encode the structure of the transition - separating sister lineages and ordering the cells along the bifurcation axis. Having established this correspondence under known ground truth, we establish that these results support our use of the term FTU in this paper.

We next turn to the analysis of EMT datasets to illustrate how scCont resolves different lineages and the context-dependency of EMT. Based on the sensitivity analysis (Fig. 2E-H and Supplementary Fig. 1), we determine that *k* = 3-10 is a suitable range for most datasets depending on the conditions. To select *k* for the EMT datasets, we swept *k* from 3 to 10 on the MCF10A TGFB1 data (Supplementary Fig. 5). We measure latent redundancy as the effective dimensionality of the latent covariance (participation ratio; Supplementary Fig. 5A), the mean pairwise Jaccard index of enriched GO terms between latent features (Supplementary Fig. 5B), and the temporal locality as the fraction of positive pairs drawn from the same timepoint (Supplementary Fig. 5C). *k* =3 gave the highest effective dimensionality, the lowest functional overlap between features, and the largest fraction of within-timepoint pairs, so we use *k* =3 for all EMT analyses. PCA projections of the latent space for *k* =3, 5, 7 and 10 (Supplementary Fig. 5D) show that lower *k* retains finer timepoint structure. Considering EMT is a continuum where multiple genes can be involved in different functional programs, we also opt for a lower *k* compared to the simulation data, where program genes are only involved in one functional group for ease of calculations.

### scCont resolves distinct temporal modules in EMT

By spatially clustering the FTUs in PCA space, scCont groups FTUs into modular transition units that exhibit coordinated temporal behavior. Figure 3 illustrates how these modular units resolve both coherent and bifurcating state modules. This clustering step identifies three distinct modules in the MCF10A dataset, each with unique GO enrichment terms. Module 0 captures a strong, temporally correlated latent state transition, isolating the FTUs that significantly drive the progressive transition. Module 1 shows a temporally uncorrelated profile, indicative of background cellular processes. Module 2 captures a bifurcating pathway, revealing divergent gene expression programs that emerge over time.

GO enrichment applied to the high-scoring genes shows that all modules are enriched at a significant level for EMT-relevant biological processes (Fig. 3). This functional annotation exposes the high-level mechanisms characterizing each of these module units: Module 0 is enriched for terms associated with aggressive migration during EMT (anchoring junctions, focal adhesion, cell motility). Conversely, Module 1 is enriched for general maintenance terms consistent with background expression (tissue development, tube morphogenesis). Lastly, Module 2 is enriched for processes explaining the temporal bifurcation in the data (metabolic processes, response to hormone).

**Figure 3.**
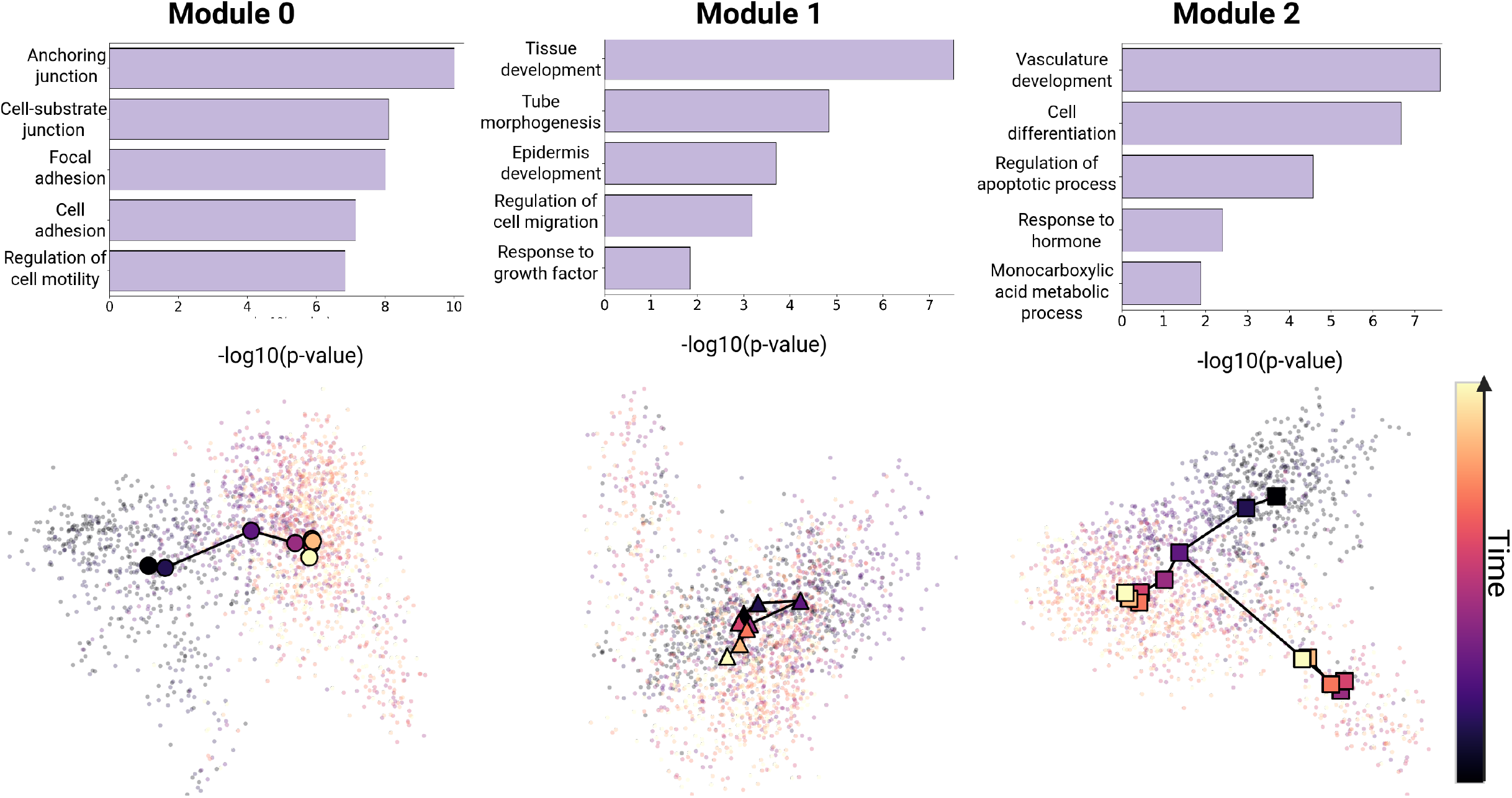
Spatial Clustering of FTUs Reveals Distinct EMT Functional Groups. **(Top) GO Enrichment:** Enrichment bar plots for each cluster are ordered in terms of statistical significance. The listed GO terms under each cluster are either processes uniquely enriched in that specific cluster, or processes where that cluster exhibits the highest comparative enrichment. **(Bottom) Context-Adaptive State Projections:** PCA plots of cells based on the aggregated functional transition units (FTUs) in each context-adaptive cluster. Cells are colored by timepoint (0-7 days) to visualize temporal progression. Modules 0, 1, and 2 comprise 11, 10, and 11 FTUs, respectively.

While the clustering of FTUs highlights broad, state-level network reconfigurations, we next perform an in-depth analysis of individual FTUs to map the functional groups driving these broader state changes. Collectively, these results demonstrate that scCont resolves distinct biological modules from continuous scRNA-seq data.

### scCont resolves core EMT genes and divergent lineage fates

To dissect the molecular drivers of these topologies, we restrict our downstream analysis to the FTUs defining the coherent (Module 0) and bifurcating (Module 2) dynamics. Figure 4 shows the PCA plots of FTUs in Modules 0 and 2. Centroids in the PCA plots are calculated by averaging the positions of cells within the respective timepoints weighted by the cells’ latent feature value. This weighted calculation means that latent state transitions gravitate to cells with higher gene expression identified by SHAP. We assign functional identities to these modular transition units via literature-based annotation of their statistically derived top-contributing genes (see Supplementary Materials for literature search results).

**Figure 4.**
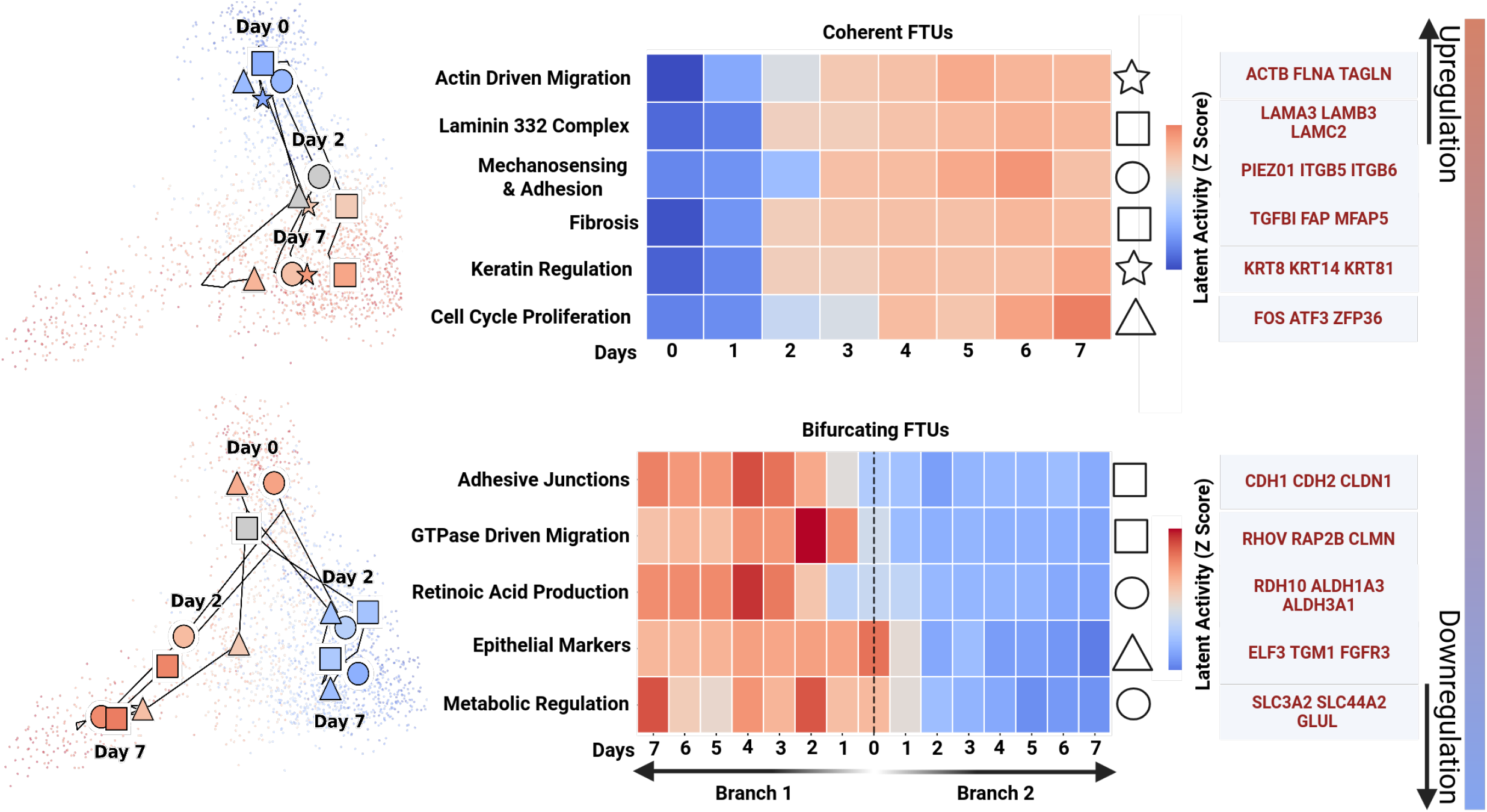
scCont Functionally Resolves Coherent vs. Bifurcating EMT Functional Transition Units (FTUs). **(Top) Coherent FTUs:** Analysis of FTUs exhibiting coordinated temporal transitions. **(Bottom) Bifurcating FTUs:** Analysis of FTUs driving the temporal divergence into distinct states. **(Left) PCA Plots:** Scatterplots visualize average FTU activation for each cell in PCA space. PCA plots are specifically highlighting FTUs from Modules 0 and 2. Feature paths connect the centroids of each timepoint (Days 0-7) with centroid coordinate calculated as the average of the PCA coordinates weighted by the FTU’s activation value. Only Days 0, 2, and 7 are explicitly labeled for clarity. **(Middle) FTU Annotation:** Symbols (square, circle, triangle) next to the heatmap correspond to the plotted FTU feature paths. Each row on the heatmap displays aggregate modular activity calculated as the sum of SHAP values from a single FTU. **(Right) Modular Gene Sets:** Gene lists shown represent examples of the statistically derived driving genes for each FTU.

The coherent cluster reveals the core EMT program-upregulated modules activated across the entire transitioning population, while the bifurcating cluster reveals a dynamic split into distinct mesenchymal states. As shown in Figure 4, we observe the enrichment of key migratory genes within the coherent FTUs, including actin- dependent cytoskeletal remodeling, mechanosensory adhesive factors, and Laminin-332 mediated cohesion, which are all fundamental processes for the E/M transition (30; 31).

However, the bifurcating FTUs exhibit phenotypic heterogeneity, representing a temporal divergence into distinct mesenchymal states. While both branches share the core migratory machinery, they differ significantly in their regulatory programming. Specifically, we identify one branch defined by an FTU enriched for retinoic acid production and epithelial genes such as ELF3 which are typically associated with epithelial maintenance (32). The global presence of aggressive, migratory factors alongside the local retention of retinoic acid production in Branch 1 suggest that the bifurcating units capture a hybrid E/M state, where cells simultaneously exhibit epithelial and mesenchymal characteristics. The concurrent upregulation of adhesion factors CDH1 and CDH2 (the cadherin switch) is further corroboration of a partial transition state (33). By decoupling the loss of epithelial identity from the gain of motility, this modular approach resolves complex, non-binary EMT outcomes.

### FTUs reveal the context-dependency of the EMT process

Next, we illustrate context-dependent analysis via scCont by applying it to 12 EMT timecourse datasets from Cook et al. (3), spanning four cell lines (A549, MCF7, DU145, OVCA420) treated with three distinct inducers: epidermal growth factor (EGF), TGFB1, and tumor necrosis factor (TNF). In these experiments, cells were treated for 8 days followed by inducer withdrawal to assess transcriptional reversibility. Previous studies establish that cells treated with EGF often retain epithelial identity, TGFB1 is a potent EMT inducer, and TNF drives an EMT inflammatory phenotype (34; 35; 36). Given the strong conservation of cell line specific signals reported in the original study (3), we focus downstream analysis on the A549 cell line across all three treatments to evaluate the cross-dataset alignment of FTUs and isolate inducer-specific adaptive reconfigurations (Figure 5). Analysis for MTUs for all other datasets can be found in Supplementary Table 1.

**Figure 5.**
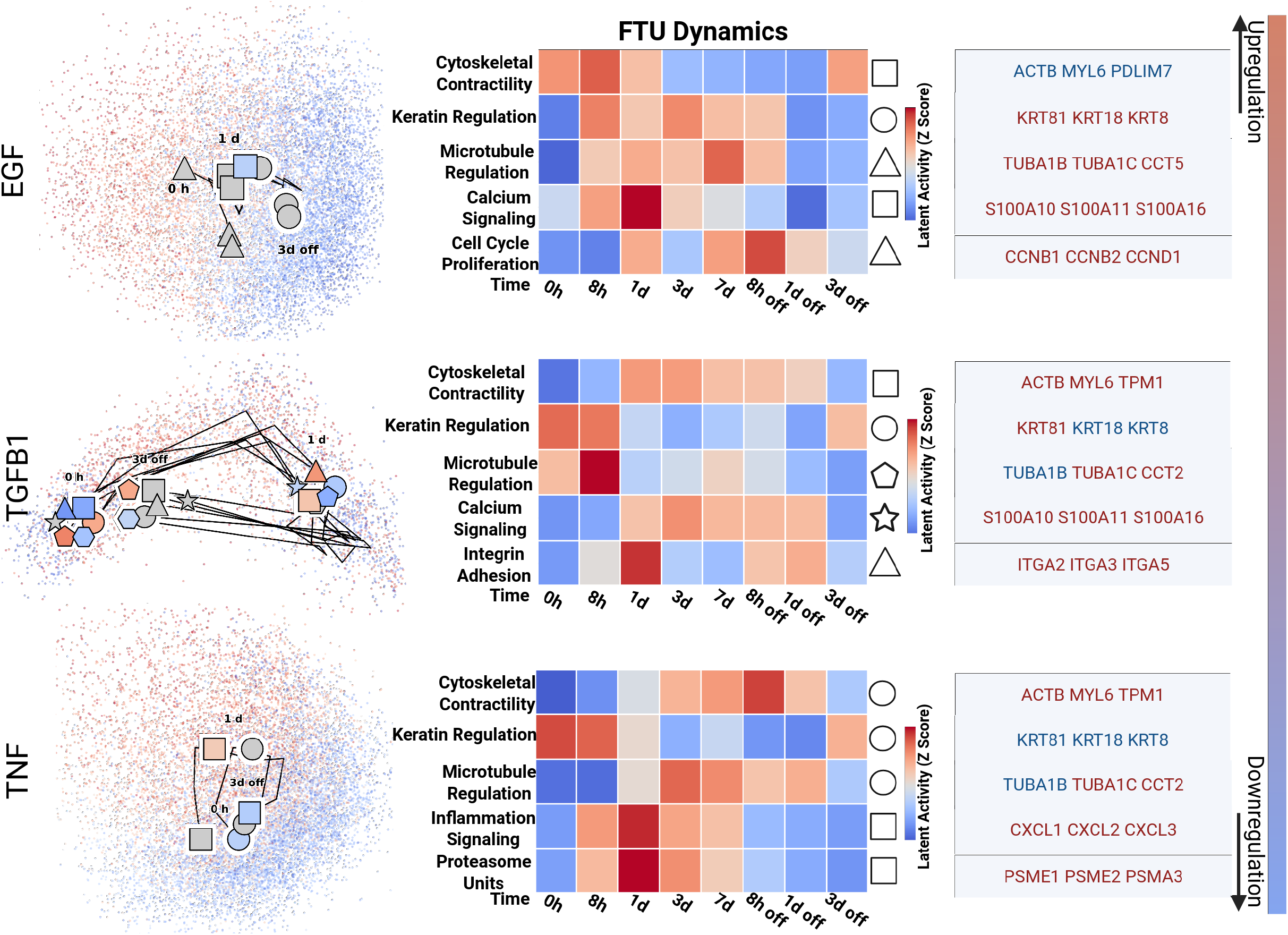
Application of scCont to A549 cells treated with EGF, TGFB1, or TNF. (Left) Functional Transition Unit Paths (FTUs): PCA projections of functional transition unit (FTU) activations showing distinct topological responses to different inducers over the treatment course. **(Center) Modular FTU Dynamics:** Heatmaps display the modular activity of identified FTUs via aggregated SHAP values. Symbols next to the heatmaps correspond to specific FTU paths in the PCA space. **(Right) Modular Gene Sets:** Examples of statistically derived driving genes defining each FTU module.

Notably, scCont identifies a core set of conserved functional modules-including cytoskeletal contractility, keratin regulation, and microtubule dynamics-across all three conditions, yet resolves three distinct, context-dependent regulatory modules. First, in the EGF condition, the extracted FTUs reveal a unique partial EMT signature: while microtubule regulation is upregulated (indicating enhanced motility), cytoskeletal contractility is downregulated, and epithelial keratins (KRT8/18) are retained (37). This configuration aligns with EGF’s role in promoting proliferation-captured by a distinct cell cycle FTU (34)-without driving a complete mesenchymal transition.

In contrast, the TGFB1 condition exhibits a complete EMT program characterized by the adaptive reconfiguration of these same FTUs. Cytoskeletal contractility and integrin adhesion modules are strongly upregulated, consistent with increases in migratory potential (34). Concurrently, epithelial markers (KRT8/18) are downregulated within the relevant FTUs, while mesenchymal- associated keratins (KRT81) and microtubule regulators (TUBA1C, CCT2) are upregulated, marking a comprehensive loss of epithelial identity (37; 38; 39; 40). Finally, the TNF dataset mirrors the migratory module activation seen with TGFB1 but is uniquely distinguished by the concurrent enrichment of inflammation and proteasome FTUs (36). The differential activation of these conserved modules across treatments demonstrates scCont’s capacity to perform cross-dataset module alignment and explicitly map how genetic context dynamically reconfigures the E/M state.

### scCont captures continuous biological and temporal features in cell-state transitions compared to other state-of-the-art models

While simulated data provides useful ground truth information to evaluate how well the models perform, we provide an additional comparison on the MCF10A TGFB1 dataset to see if the performance of the models transfers to real applications. Because we lack ground truth functional programs, we use GO term enrichment on each latent feature to see if individual latent features are capable of capturing biologically coherent functional groups. We also measure temporal structure using known timepoint labels from the datset. To ensure a consistent comparison, all models are parameterized with 32 latent features just like in the comparison for the simulated data. LDVAE was parameterized with 20 latent features in its original study (22), and other models that use NMF for scRNA data analysis also confirm that 32 latent features is a suitable parameter for NMF (28; 29).

Figure 6 evaluates these five models across temporal and biological recovery metrics on the MCF10A TGFB1 dataset and demonstrates a strict trade-off between temporal manifold preservation and functional interpretability in baseline models. While scVI, LDVAE, and scCont successfully preserve global temporal ordering, achieving the highest predictive power, NMF substantially degrades the continuous temporal information. Interestingly, compared to Figure 2L, all models perform well for timepoint prediction except NMF on the real data, while scCont performed as well or better than all other methods. One explanation is that with the simulated data, we have access to continuous ground truth pseudotime information, while for the MCF10A TGFB1 data, we only have access to discrete timepoints. The prediction of discrete timepoints could be an easier task for the models compared to the continuous pseudotime prediction, leading to an improvement in baseline model performance. To continue, we evaluate the latent features of each model by measuring GO term enrichment for each method. In Figure 6B, we first measure the proportion of latent features enriched for any GO term. We notably observe that LDVAE consistently lags behind all other methods in terms of identifying coherent functional groups. Next, we test for the extraction of specific EMT FTU enrichment across five random initializations (Fig. 6C). Many researchers conclude that the known hallmarks of EMT include the loss of apical-basal polarity, the reduction of cell-cell adhesion via tight junctions, the acquisition of focal adhesions, the increased motility and invasive ability, and the degradation of the extracellular matrix (41; 42; 43). Therefore, we specifically measure the enrichment for GO terms shown in Figure 6C and their child terms. Overall, we observe LDVAE and scVI fail to consistently isolate modular biology from this dataset compared to scCont and NMF. To compare these results to the simulated dataset, we again notice a difference between how the models performed. LDVAE performed well on program recovery in Figure 2I, but does not capture coherent functional groups via GO term enrichment analysis in Figure 6B-C. This discrepancy could again be attributed to no one gene acting in more than one program in the simulated data, meaning that measurement of a functional group in a real dataset could be a harder task compared to the simulated data since gene sets can be overlapping.

**Figure 6.**
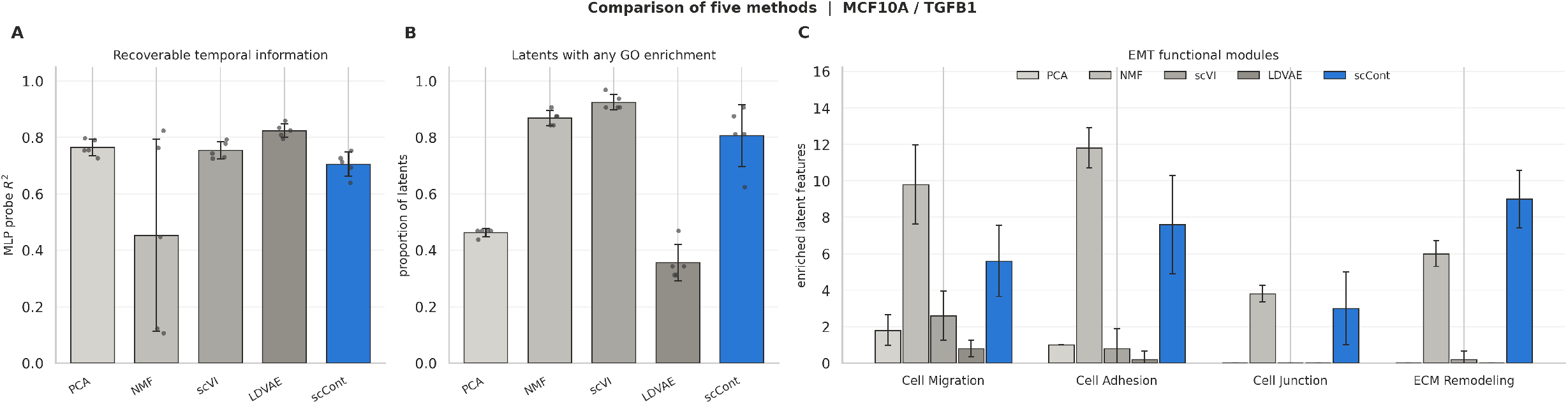
Benchmarking scCont against state-of-the-art models. **A) Recoverable Temporal Information:** Cross-validation study assessing the *R*^2^ predictive power of each model. **B) Latent Space Utilization across Random Seeds:** The proportion of enriched latent features is reported across five random initializations. **C) EMT Functional Transition Units (FTUs) across Random Initializations:** The number of enriched EMT related FTUs for each model across five random seeds. GO enrichment was performed to find listed terms and their child terms (*adj. p-val <* 0.05). All benchmarking experiments in this figure use the MCF10A TGFB1 dataset.

Supplementary Figure 6 provides visual confirmation of these representational constraints. The latent features of NMF and LDVAE, in particular, show a rigid, spatial compartmentalization of the input space consistent with linear models (44; 16). This compartmentalization does not capture the continuous information in timecourse scRNA-seq data, where continuous cell lineage fates must be preserved. In contrast, the compositional architecture of scCont resolves this dichotomy: by combining a *k* -NN-based contrastive objective to explicitly preserve temporal topology with a dedicated interpretability stack, scCont accurately maps these lineages in the input space, extracting modular transition units without compromising the chronological biological progression. Therefore, we conclude that on simulated and real datasets, scCont is a top performer in most measures of performance. We establish in Figure 2 that with known functional programs, scCont recovers these programs, while preserving temporal information and disentangling the true signal from the background genes. We, then, show that these attributes extend to a real dataset, while some other baseline models fail to generalize.

## Discussion

In this paper, we present scCont, an unsupervised, compositional contrastive learning framework that combines the predictive power of neural networks with the biological interpretability required for single-cell analysis. While deep learning has advanced omics data processing, its application has often been hindered by the opaque nature of these models. scCont’s primary innovation lies in its compositional architecture: using a *k* -NN-based positive pair selection method to explicitly preserve temporal topology and encapsulate functional programs in an unsupervised manner. Because of this algorithmic integration, scCont effectively decouples complex transcriptional signals into context-adaptive FTUs, offering a transparent view into the molecular drivers of EMT. Our application of scCont to timecourse data reveals topological structures that standard linear methods could not fully resolve. Specifically, scCont identified a bifurcation in the EMT continuum, distinguishing a conserved coherent program driven by cytoskeletal remodeling from a divergent lineage characterized by retinoic acid signaling and retention of epithelial markers. This ability to capture branching topologies and hybrid E/M states (upregulation of mesenchymal factors with the retention of epithelial characteristics) highlights the necessity of nonlinear approaches for modeling developmental plasticity and adaptive state networks.

When we evaluate scCont against four other baseline models across simulated and real datasets, we establish that scCont is a top model for all measures of performance. Additionally, we illustrate that scCont is capable of identifying ground truth functional programs and encapsulating pseudotime information relevant to ground truth lineages. These results transfer well when the model is evaluated on the MCF10A TGFB1 data. As a result, scCont effectively encompasses program information while recovering temporal resolution when compared to other models.

Despite these advantages, the compositional nature of scCont presents distinct challenges. Algorithmically, the model’s success relies on the assumption that the initial *k* -NN graph accurately captures the true biological progression. Furthermore, unlike PCA or NMF, which are computationally efficient and run on standard hardware, scCont performs better with GPU acceleration for optimal performance and scales less efficiently with dataset size. Similarly, while the SHAP package provides non-linear feature attribution, its computational overhead scales linearly with the number of cells, creating a computational bottleneck for atlas-scale datasets (26). Collectively, these constraints represent a classic trade-off between computational cost and model expressivity.

Looking forward, the biologically-disentangled FTUs learned by scCont offer a robust foundation for unsupervised trajectory inference. Applying methods such as Monocle3 to scCont’s modular latent space could enable fully unsupervised trajectory construction of functional gene programs in novel biological systems. Additionally, the foundation of scCont - the positive pair selection method - can serve as a powerful tool for other neural network methods that aim to learn some underlying structure in data in an unsupervised manner. Ultimately, we present scCont as a robust, open-source analytical tool for the genomics community. scCont moves interpretable deep learning toward practical application, providing researchers with a framework to examine the internal representations of neural networks. By explicitly modeling temporal manifolds, scCont empowers the field to extract not just mathematical abstractions, but modular, tangible, context-specific biological mechanisms from complex single-cell datasets.

## Methods

### Contrastive Learning Framework

#### Network Architecture

The scCont framework was implemented using PyTorch. The encoder function *f(•)* was designed as a four-layer MLP mapping high-dimensional gene expression to a 32-dimensional latent space. The layer dimensions were set to 3000 (input genes) *→* 2048 *→* 1024 *→* 64 *→* 32. Rectified Linear Unit (ReLU) and Batch Normalization were applied between layers. The projector function *g(•)* consisted of a two-layer MLP mapping the 32-dimensional latent representation to a 16-dimensional output space for contrastive loss calculation.

#### Training Strategy

Positive pairs for contrastive training were generated via *k* -NN graph selection in the normalized gene expression space (See Supp. Fig. 4A and 5C for additional results). The model was trained using the InfoNCE loss function with a temperature of *τ* = 0.20. Optimization and learning were performed using the Adam optimizer with a learning rate of 1e-4. Following the framework from Chen et al., we employed an iteration-based training strategy, processing 20,000 batches of size 256 (10). This computational budget ensured model convergence across all datasets.

### Interpretability & Downstream Analysis

#### Network Attribution

To determine the biological features driving the latent representations, we employed the DeepSHAP explainer from the SHapley Additive exPlanations (SHAP) Python package (26). A reference distribution of 100 cells was used for each instance in which SHAP was applied. We calculated the mean absolute SHAP value for each gene across all cells to derive global feature importance for each latent dimension. Functional annotation was performed using g:Profiler to map top-contributing genes (*z >* 2.57, *p <* 0.01) to Gene Ontology (GO) biological processes (*adj. p-val <* 0.05). To understand how well SHAP scales with data, a scalability test for SHAP was conducted. SHAP’s runtime increases linearly with more data with an average of 170 ms/cell for all runs conducted.

#### Spatial Clustering of FTUs

To identify spatially co-varying functional modules, FTU activations were projected onto the original PCA space on a 40x40 grid, and the activation of each FTU was aggregated within each bin. This resulted in a 2D density map for each FTU. These density maps were flattened into feature vectors, and pairwise distances were calculated using the correlation distance metric (1 - *r*). We then applied agglomerative hierarchical clustering with average linkage on the distance matrix to partition the FTUs into distinct spatial groups. For each dataset, an elbow plot was generated to determine the optimal number of clusters. This clustering method was used to overcome single-cell noise/sparsity and allow downstream analysis to focus on similar expression patterns identified by the FTUs.

### Simulated Datasets

#### Dataset Generation

SERGIO simulates expression for every gene in every cell by numerically integrating a stochastic model of transcription in which each gene’s production rate is determined by the current concentrations of its regulators and the interaction strengths specified in the inputs. Cells of a given state are drawn from the resulting distribution; in differentiation mode each state is simulated downstream of its parent, so cells are sampled along continuous trajectories rather than from discrete clusters. We then applied SERGIO’s technical-noise model, which adds expression outliers, cell-to-cell variation in library size and dropout before conversion to UMI counts. The output is a count matrix over the program-associated genes.

#### Gene Regulatory Network Construction

Each simulated dataset is generated from a directed acyclic graph (DAG) comprising either 16 or 32 disjoint sub-networks with each disjoint sub-network representing a gene program. Each gene program has one master regulator with target genes partitioned into two layers: 75% of target genes are regulated by the master regulator and the remaining 25% are regulated by a direct target from the master regulator, meaning that all these genes are within two hops away from the master regulator. No edges cross between programs, so every gene is only reachable from one master regulator. This entire process makes up either 400 or 800 program genes in the simulation.

To reach a realistic feature count that could distract the models from the actual functional gene programs, each background gene is a copy of a uniformly randomly chosen simulated gene whose values are permuted across cells. This process preserves the gene count distribution and sequencing depth, which allows the simulation to mimic other noisy genes that may have passed a highly variable gene filtering step. However, the random permutation across cells destroys the program structure associated with the gene. The result is a set of background genes that are within the proper distribution of the functional genes created through SERGIO but have no inherent program structure or relationship with other genes.

#### Pseudotime Construction

Ground-truth pseudotime is derived directly from the SERGIO dynamic simulations. SERGIO simulates each lineage state as a set if trajectories evolving from a parent state toward a new steady state, generating a time step for each cell. The index of each cell is used to calculate its pseudotime value. For a cell sampled at step *t* from lineage state *b* with trajectory length *T*_*b*_, pseudotime is defined as *depth*(*b*) + (*T*_*b*_ *™* 1), where *depth*(*b*) is the distance of state *b* from the root of the lineage tree. The result is then scaled to a range between 0 and 1. This formulation produces a pseudotime value that increases continuously along each root-to-leaf path.

### Data Processing

#### MCF10A TGFB1 Dataset

The MCF10A dataset was acquired from the Gene Expression Omnibus (GEO Accession: GSE200981). Data processing was performed in Python using ScanPy and included: 1) filtering low-quality cells (*<* 200 genes expressed); 2) filtering genes expressed in fewer than 3 cells; 3) doublet detection and removal via Scrublet; 4) library size normalization (10,000 counts per cell) and log-transformation; 5) selection of the top 3,000 highly variable genes; 6) regressing out cell cycle scores, mitochondrial percentage, and total counts; and 7) scaling data to unit variance.

#### Cook et al. Datasets

Pre-processed Seurat objects for the A549, MCF7, DU145, and OVCA420 cell lines were obtained from the original authors (3). These datasets were converted to AnnData format and, as they followed an identical preprocessing pipeline to the MCF10A dataset described above, were used without further modification. All models were trained using a fixed random seed to ensure reproducibility. For all EMT datasets, *k* =3 was used for scCont.

### Benchmarking

scCont was compared against LDVAE, scVI, NMF, and PCA. Across all models, the identical set of 3,000 highly variable genes (HVGs) was used with input data tailored to satisfy each architecture’s specific requirements. Because scVI and LDVAE require raw, unnormalized counts, no additional processing was performed beyond HVG selection. For NMF, which requires non-negative inputs, the data underwent the standard pre-processing pipeline, excluding data scaling and the regression of cell cycle and mitochondrial signals to preserve non-negativity. PCA and scCont used scaled, normalized data. All models were initialized with 32 latent space dimensions, a choice validated by existing literature as suitable across these architectures (20; 22; 28; 29). All benchmarking comparisons were conducted using the MCF10A TGFB1 dataset across the same five random seeds. To ensure consistency during Gene Ontology (GO) enrichment analysis and simulated dataset analysis, the top 100 genes from each latent feature were tested for every model. We provide runtime analysis on all five methods in Supplementary Figure 3.

## Abbreviations

(scRNA): single-cell RNA
(NMF): non-negative matrix factorization
(FTUs): functional transition units
(MTUs): modular transition units
(EMT): epithelial-to-mesenchymal transition
(SHAP): Shapley Additive exPlanations
(LDVAE): linearly decoded variational autoencoder

## Data Availability

All source code, processed datasets, GO enrichment tables, and trained models required to reproduce the results are available on the publicly available scCont GitHub page at https://github.com/nramani611/scCont. Supplementary Materials include the functional annotation citations. A Jupyter notebook tutorial demonstrating the analysis pipeline is also provided in the repository. All scripts used to generate the SERGIO datasets are also found on the GitHub page.

## Key Messages

- Deep understanding of cellular state transitions requires continuous tracking of functional gene programs.
- We present scCont, which uniquely provides unsupervised temporal tracking of functional modules of cellular state transitions.
- scCont’s technical innovation relies on temporally informative pair selection for contrastive learning combined with strong interpretability methods.

---

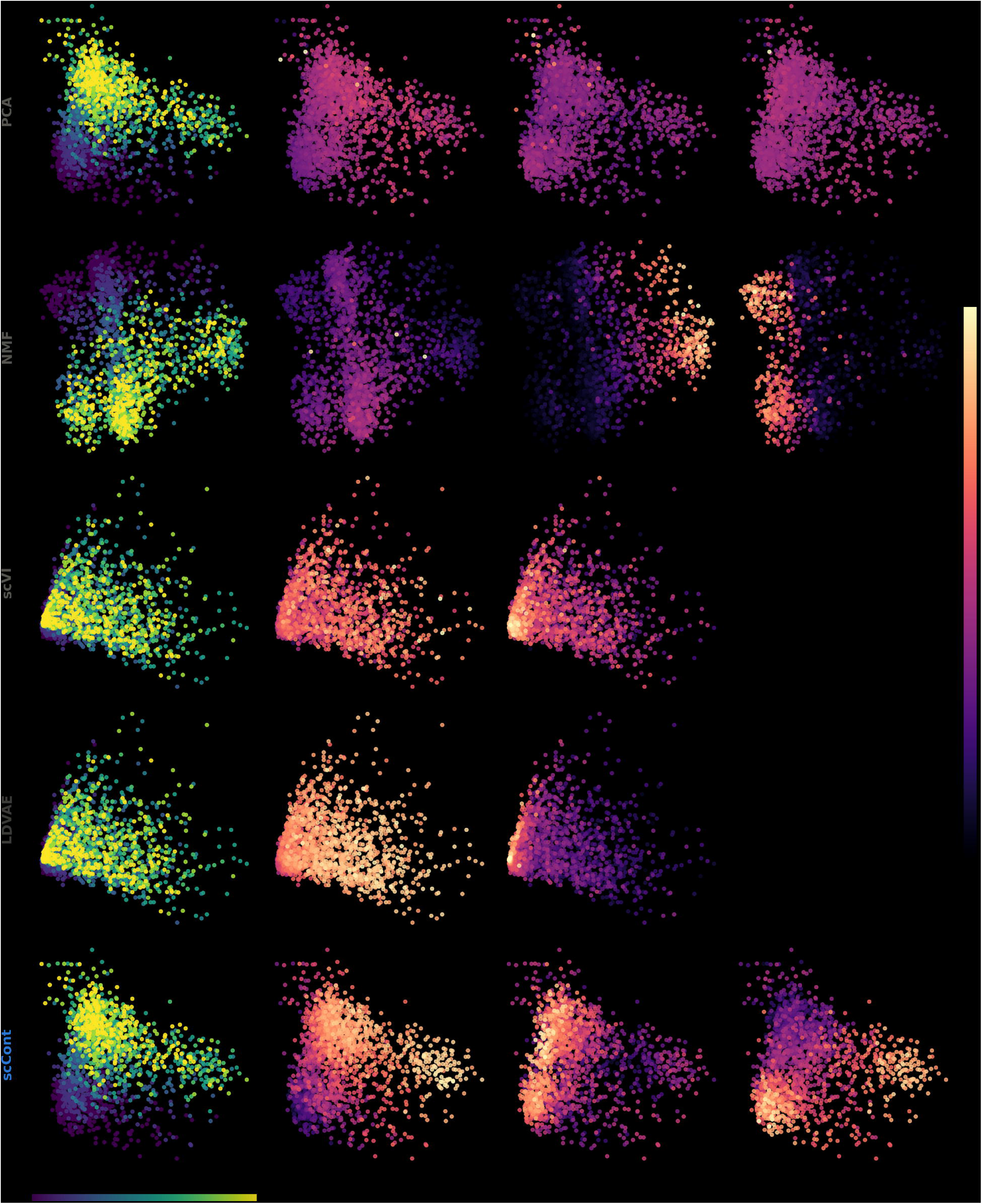

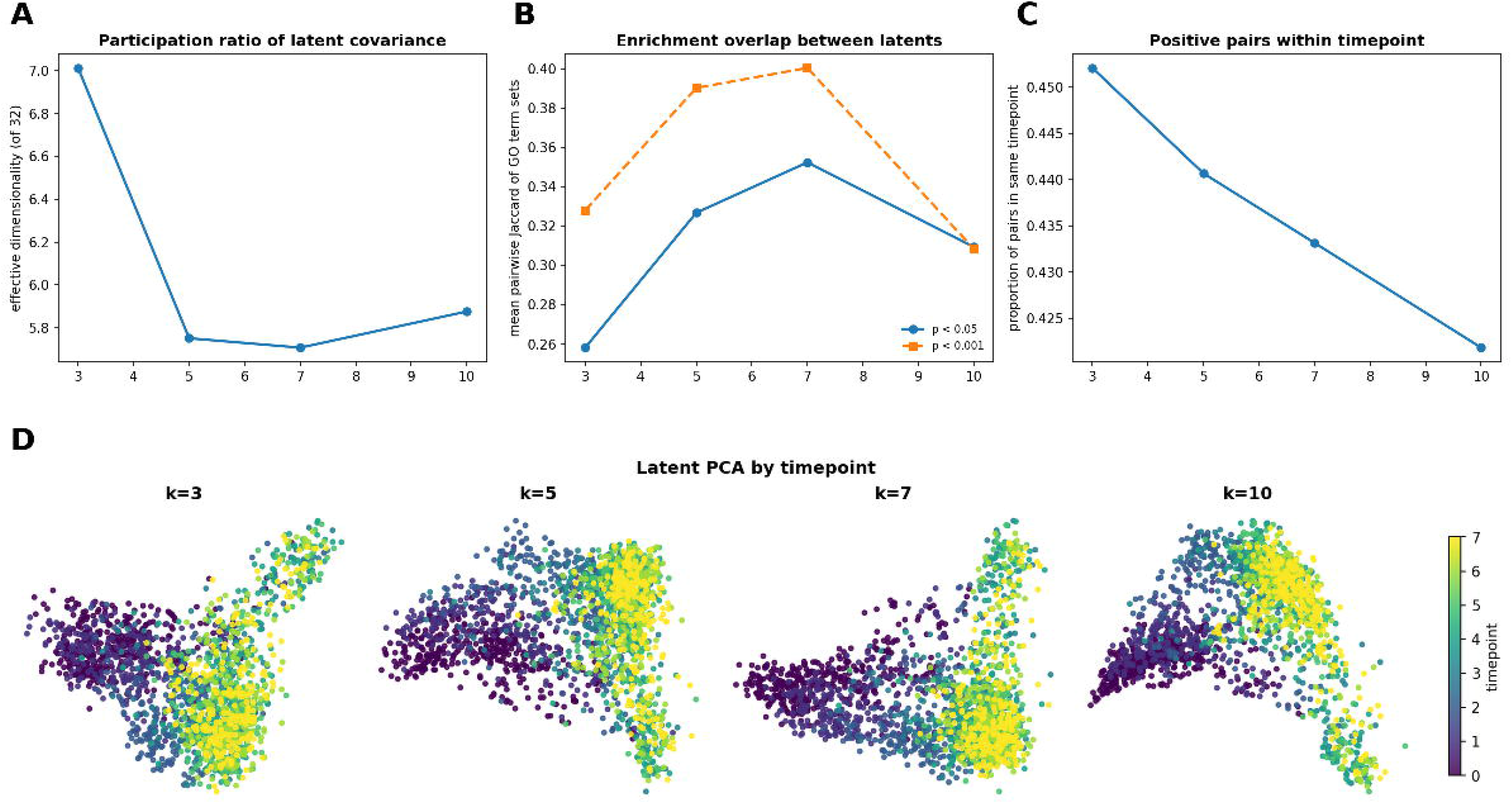

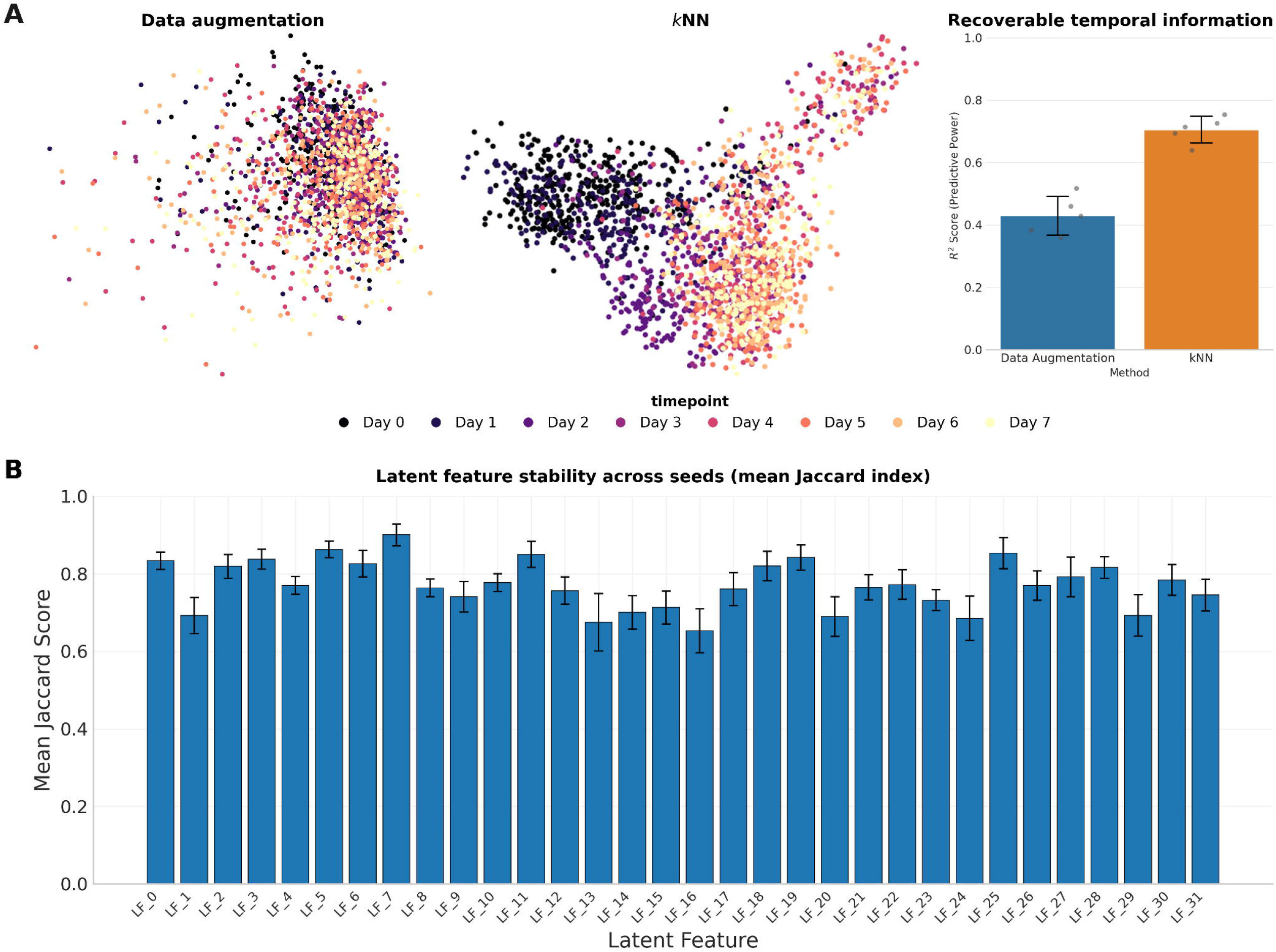

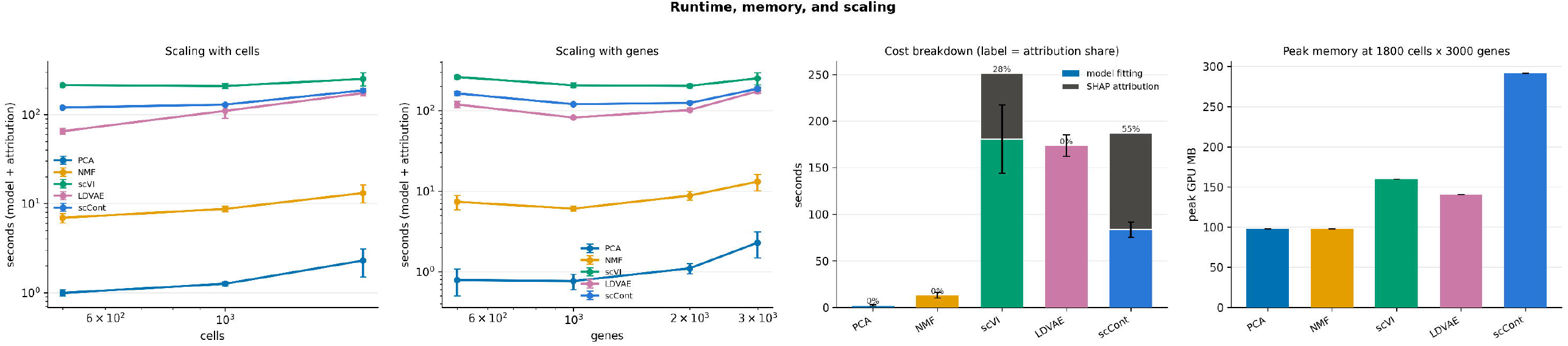

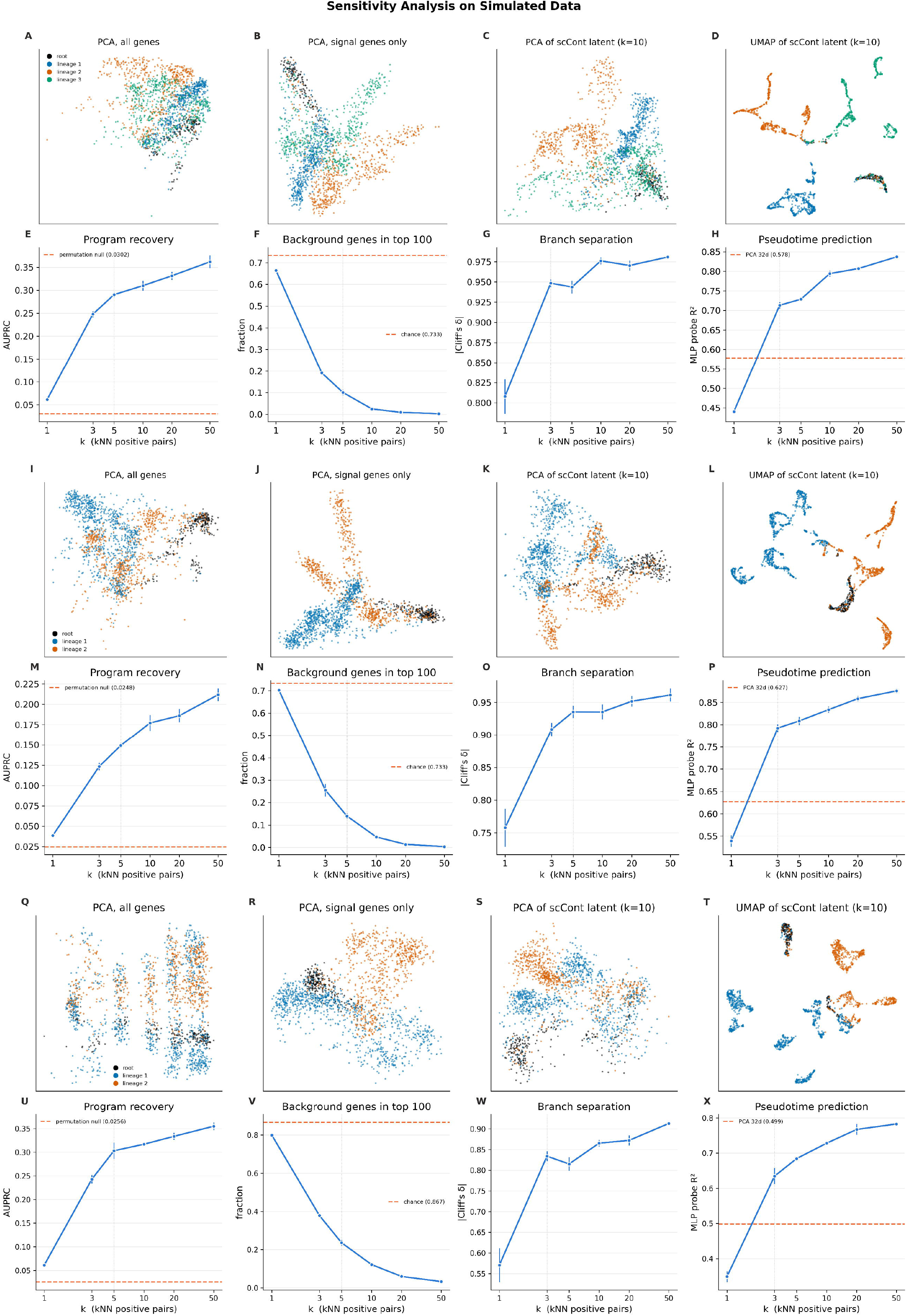

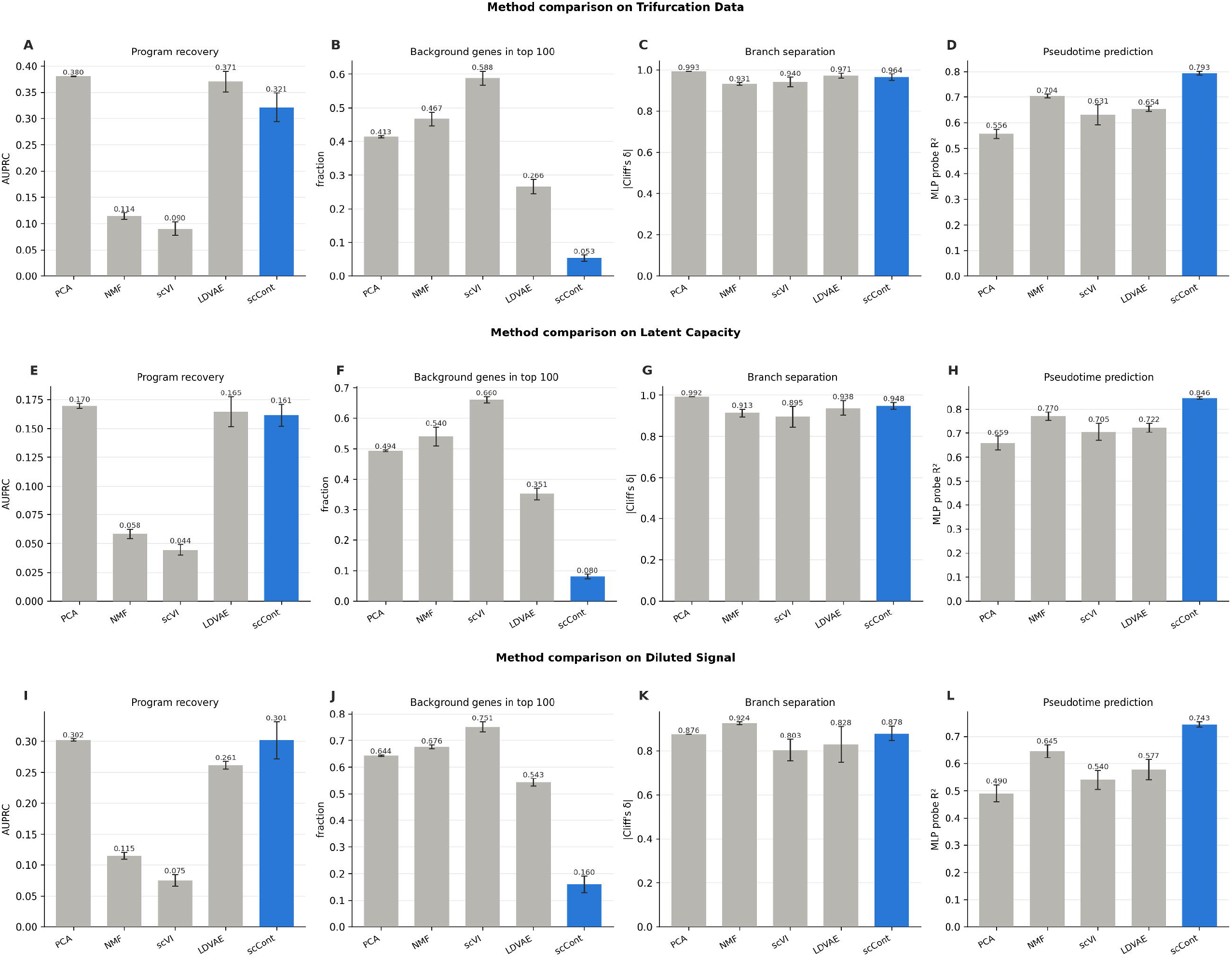

## References

1. Zhang CX, Huang RYJ, Sheng G, Thiery JP. Epithelial-mesenchymal transition. Cell. 2025;188(20):5436–86.

2. Jordan NV, Johnson GL, Abell AN. Tracking the intermediate stages of epithelial-mesenchymal transition in epithelial stem cells and cancer. Cell Cycle. 2011;10(17):2865–73.

3. Cook DP, Vanderhyden BC. Context specificity of the EMT transcriptional response. Nature Communications. 2020;11(1):2142.

4. Sun S, Zhu J, Ma Y, Zhou X. Accuracy, robustness and scalability of dimensionality reduction methods for single-cell RNA-seq analysis. Genome Biology. 2019;20(1):269.

5. Kotliar D, Veres A, Nagy MA, Tabrizi S, Hodis E, Melton DA, et al. Identifying gene expression programs of cell-type identity and cellular activity with single-cell RNA-Seq. eLife. 2019 jul;8:e43803. Available from: 10.7554/eLife.43803.

6. Stuart T, Butler A, Hoffman P, Hafemeister C, Papalexi E, Mauck WM, et al. Comprehensive Integration of Single-Cell Data. Cell. 2019;177(7):1888–902.e21.

7. Li S, Hua H, Chen S. Graph neural networks for single-cell omics data: a review of approaches and applications. Briefings in Bioinformatics. 2025;26(2):bbaf109.

8. Fan FL, Xiong J, Li M, Wang G. On Interpretability of Artificial Neural Networks: A Survey. IEEE Transactions on Radiation and Plasma Medical Sciences. 2021;5(6):741–60.

9. Singh R, Sledzieski S, Bryson B, Cowen L, Berger B. Contrastive learning in protein language space predicts interactions between drugs and protein targets. Proceedings of the National Academy of Sciences. 2023;120(24):e2220778120.

10. Chen T, Kornblith S, Norouzi M, Hinton G. A Simple Framework for Contrastive Learning of Visual Representations. arXiv. 2020.

11. Oord Avd, Li Y, Vinyals O. Representation Learning with Contrastive Predictive Coding. arXiv. 2018.

12. Han W, Cheng Y, Chen J, Zhong H, Hu Z, Chen S, et al. Self-supervised contrastive learning for integrative single cell RNA-seq data analysis. Briefings in Bioinformatics. 2022 09;23(5):bbac377. Available from: 10.1093/bib/bbac377.

13. Yang M, Yang Y, Xie C, Ni M, Liu J, Yang H, et al. Contrastive learning enables rapid mapping to multimodal single-cell atlas of multimillion scale. Nature Machine Intelligence. 2022;4(8):696–709.

14. Dibaeinia P, Sinha S. SERGIO: A Single-Cell Expression Simulator Guided by Gene Regulatory Networks. Cell Systems. 2020;11(3):252–71.e11.

15. Paul I, Bolzan D, Youssef A, Gagnon KA, Hook H, Karemore G, et al. Parallelized multidimensional analytic framework applied to mammary epithelial cells uncovers regulatory principles in EMT. Nature Communications. 2023;14(1):688.

16. Stein-O’Brien GL, Arora R, Culhane AC, Favorov AV, Garmire LX, Greene CS, et al. Enter the Matrix: Factorization Uncovers Knowledge from Omics. Trends in Genetics. 2018;34(10):790–805.

17. Hozumi Y, Wei GW. Analyzing single cell RNA sequencing with topological nonnegative matrix factorization. Journal of Computational and Applied Mathematics. 2024;445:115842. Available from: https://www.sciencedirect.com/science/article/pii/S0377042724000918.

18. Shao C, Höfer T. Robust classification of single-cell transcriptome data by nonnegative matrix factorization. Bioinformatics. 2016;33(2):235–42.

19. Zheng X, Jin S, Nie Q, Zou X. scRCMF: Identification of Cell Subpopulations and Transition States From Single-Cell Transcriptomes. IEEE Transactions on Biomedical Engineering. 2018;67(5):1418–28.

20. Lopez R, Regier J, Cole MB, Jordan MI, Yosef N. Deep generative modeling for single-cell transcriptomics. Nature Methods. 2018;15(12):1053–8.

21. Lotfollahi M, Wolf FA, Theis FJ. scGen predicts single-cell perturbation responses. Nature Methods. 2019;16(8):715–21.

22. Svensson V, Gayoso A, Yosef N, Pachter L. Interpretable factor models of single-cell RNA-seq via variational autoencoders. Bioinformatics. 2020 03;36(11):3418–21. Available from: 10.1093/bioinformatics/btaa169.

23. Li WY, Murtaza G, Singh R. scContrast: A contrastive learning based approach for encoding single-cell gene expression data. bioRxiv. 2025. Available from: https://www.biorxiv.org/content/early/2025/04/14/2025.04.07.647292.

24. Shi Y, Wan J, Zhang X, Liang T, Yin Y. scCRT: a contrastive-based dimensionality reduction model for scRNA-seq trajectory inference. Briefings in Bioinformatics. 2024 05;25(3):bbae204. Available from: 10.1093/bib/bbae204.

25. Xiong Z, Luo J, Shi W, Liu Y, Xu Z, Wang B. scGCL: an imputation method for scRNA-seq data based on graph contrastive learning. Bioinformatics. 2023 02;39(3):btad098. Available from: 10.1093/bioinformatics/btad098.

26. Lundberg S, Lee SI. A Unified Approach to Interpreting Model Predictions. arXiv. 2017.

27. Ponce-Bobadilla AV, Schmitt V, Maier CS, Mensing S, Stodtmann S. Practical guide to SHAP analysis: Explaining supervised machine learning model predictions in drug development. Clinical and Translational Science. 2024;17(11):e70056.

28. DeBruine ZJ, Pospisilik JA, Triche TJ. Fast and interpretable non-negative matrix factorization for atlas-scale single cell data. bioRxiv. 2024:2021.09.01.458620.

29. Palla G, Ferrero E. Latent Factor Modeling of scRNA-Seq Data Uncovers Dysregulated Pathways in Autoimmune Disease Patients. iScience. 2020;23(9):101451.

30. Gjorevski N, Boghaert E, Nelson CM. Regulation of Epithelial-Mesenchymal Transition by Transmission of Mechanical Stress through Epithelial Tissues. Cancer Microenvironment. 2012;5(1):29–38.

31. Marinkovich MP. Laminin 332 in squamous-cell carcinoma. Nature Reviews Cancer. 2007;7(5):370–80.

32. Subbalakshmi AR, Sahoo S, Manjunatha P, Goyal S, Kasiviswanathan VA, Mahesh Y, et al. The ELF3 transcription factor is associated with an epithelial phenotype and represses epithelial-mesenchymal transition. Journal of Biological Engineering. 2023;17(1):17.

33. George JT, Jolly MK, Xu J, Somarelli J, Levine H. Survival outcomes in cancer patients predicted by a partial EMT gene expression scoring metric. Cancer Research. 2017;77(22):canres.3521.2016.

34. Buonato JM, Lan IS, Lazzara MJ. EGF augments TGFB-induced epithelial–mesenchymal transition by promoting SHP2 binding to GAB1. Journal of Cell Science. 2015;128(21):3898–909.

35. Willis BC, Borok Z. TGF-B-induced EMT: mechanisms and implications for fibrotic lung disease. American Journal of Physiology-Lung Cellular and Molecular Physiology. 2007;293(3):L525–34. PMID: 17631612. Available from: 10.1152/ajplung.00163.2007.

36. Bates RC, Mercurio AM. Tumor Necrosis Factor-a Stimulates the Epithelial-to-Mesenchymal Transition of Human Colonic Organoids. Molecular Biology of the Cell. 2003;14(5):1790–800. PMID: 12802055. Available from: 10.1091/mbc.e02-09-0583.

37. Fortier AM, Asselin E, Cadrin M. Keratin 8 and 18 Loss in Epithelial Cancer Cells Increases Collective Cell Migration and Cisplatin Sensitivity through Claudin1 Up-regulation*. Journal of Biological Chemistry. 2013;288(16):11555–71.

38. Zhang K, Liang Y, Zhang W, Zeng N, Tang S, Tian R. KRT81 Knockdown Inhibits Malignant Progression of Melanoma Through Regulating Interleukin-8. DNA and Cell Biology. 2021;40(10):1290–7. PMID: 34591651. Available from: https://journals.sagepub.com/doi/abs/10.1089/dna.2021.0317.

39. Ghozlan H, Showalter A, Lee E, Zhu X, Khaled A. Chaperonin-Containing TCP1 Complex (CCT) Promotes Breast Cancer Growth Through Correlations With Key Cell Cycle Regulators. Frontiers in Oncology. 2021 04;11:663877.

40. Li J, Chen M, Tong M, Cao Q. TUBA1C orchestrates the immunosuppressive tumor microenvironment and resistance to immune checkpoint blockade in clear cell renal cell carcinoma. Frontiers in Immunology. 2024;15:1457691.

41. Yang J, Antin P, Berx G, Blanpain C, Brabletz T, Bronner M, et al. Guidelines and definitions for research on epithelial–mesenchymal transition. Nature Reviews Molecular Cell Biology. 2020;21(6):341–52.

42. Lamouille S, Xu J, Derynck R. Molecular mechanisms of epithelial–mesenchymal transition. Nature Reviews Molecular Cell Biology. 2014;15(3):178–96.

43. Kalluri R, Weinberg RA. The basics of epithelial-mesenchymal transition. Journal of Clinical Investigation. 2009;119(6):1420–8.

44. Lee DD, Seung HS. Learning the parts of objects by non-negative matrix factorization. Nature. 1999;401(6755):788–91.

