## Supplementary Materials for "scCont: Interpretable Contrastive Learning for Functional Characterization of Cellular State Transitions"

### Additional SERGIO Simulation Runs

Supplementary Figures 1 and 2 are additional analyses of SERGIO simulated datasets. We use these datasets to see if the results are consistent across different simulation conditions. The three additional datasets test the five models with a trifurcation (800 functional genes, 16 functional programs, 2200 background genes), at maximum latent capacity (800 functional genes, 32 functional programs, 2200 background genes), and with a diluted signal (400 functional genes, 16 functional programs, 2600 background genes). Supplementary Figure 3 details a runtime analysis of all five models.

### Functional Annotation of Latent Features

We provide a detailed literature-based annotation for the top-contributing genes identifying the latent features in the MCF10A TGFB1 dataset (main text Figure 4).

#### MCF10A TGFB1 functional programs

##### *Coherent Functional Transition Units:*

Actin Driven Migration: Upregulation of ACTB, FLNA, and TAGLN drives the reorganization of the actin cytoskeleton, a prerequisite for mesenchymal motility. FLNA (Filamin A) crosslinks actin filaments to support membrane ruffling, while TAGLN (Transgelin) is an actin-bundling protein that stabilizes stress fibers necessary for cell invasion (12; 31; 28).

Laminin-332 Complex: LAMA3, LAMB3, and LAMC2 encode the subunits of the heterotrimeric Laminin-332 complex. This complex is a key component of the basement membrane that, when upregulated, promotes cell migration and tumorigenesis via interaction with integrin receptors (23).

Mechanosensing & Adhesion: PIEZO1 encodes a mechanosensitive ion channel that converts mechanical stress into biochemical signals, regulating cell stiffness and adhesion during EMT (35). ITGB5 and ITGB6 are integrin beta subunits that mediate cell-matrix interactions, critical for the activation of latent TGFB1 and the progression of fibrosis (37).

Fibrosis: FAP (Fibroblast Activation Protein), MFAP5, and TGFBI (TGF-Beta Induced) are canonical markers of myofibroblast differentiation and extracellular matrix (ECM) remodeling, indicating a fibrotic-like mesenchymal state (10; 4; 32).

Keratin Regulation: Keratins are intermediate filaments that undergo distinct "switch" patterns during EMT. While epithelial keratins are typically downregulated, the specific regulation of KRT8, KRT15, and KRT81 highlights the remodeling of the intermediate filament network required to accommodate increased cellular flexibility (18).

##### *Bifurcating Functional Transition Units:*

Adhesive Junctions: CDH1 (E-Cadherin), CDH2 (N-Cadherin), and GJA5 (Connexin 40) regulate cell-cell junctions. The simultaneous high importance of both epithelial (CDH1) and mesenchymal (CDH2) markers in this group strongly suggests a hybrid E/M state or partial EMT, where cells retain intercellular adhesion while acquiring motility traits (11; 14).

GTPase Driven Migration: RHOV and RAP2B are Rho-family GTPases that regulate cytoskeletal dynamics. RHOV is known to promote neural crest migration, while RAP2B mediates cell scattering. CLMN (Calmin) modulates Rho-GTPase activity to fine-tune adhesion dynamics during migration (7; 6; 26).

Retinoic Acid Biosynthesis & Signaling: RDH10, ALDH1A3, and ALDH3A1 are key enzymes in the retinoic acid (RA) synthesis pathway. RA signaling is a well-established inhibitor of EMT that promotes differentiation and epithelial maintenance. The upregulation of these genes in the bifurcating lineage supports the hypothesis of an EMT-resistant or alternative differentiation trajectory (1; 9; 22).

Epithelial Differentiation Drivers: ELF3 (ESE-1) is an epithelial-specific transcription factor known to repress EMT and maintain barrier function. TGM1 and FGFR3 are similarly associated with epithelial differentiation and homeostasis. Their persistence in this latent feature further corroborates the maintenance of an epithelial identity in the resistant branch (27; 16; 25).

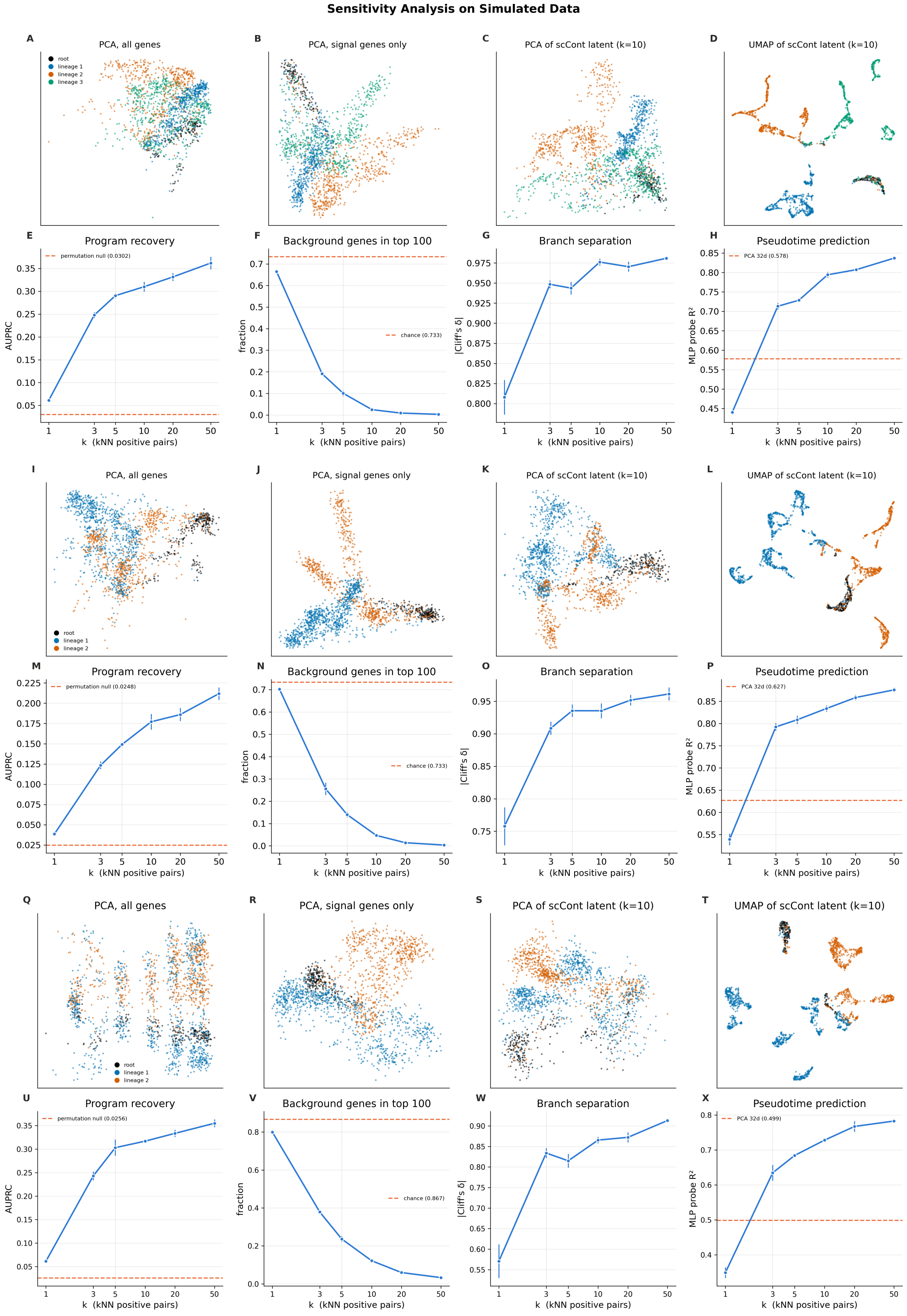


**Figure 1 scCont sensitivity analysis using three different SERGIO simulated datasets: Panels A-H)** Simulated dataset with trifurcation (3 branches) data. 800 functional genes, 16 functional programs (50 genes per functional program), and 2200 background genes. **Panels I-P) Simulated dataset on Maximum Latent Capacity.** Simulated dataset with bifurcation with latent features at maximum capacity. 800 functional genes, 32 functional programs (25 genes per functional program), and 2200 background genes. **Panels Q-X)** Simulated dataset with diluted signal. 400 functional genes, 16 functional program (25 genes per program), and 2600 background genes.

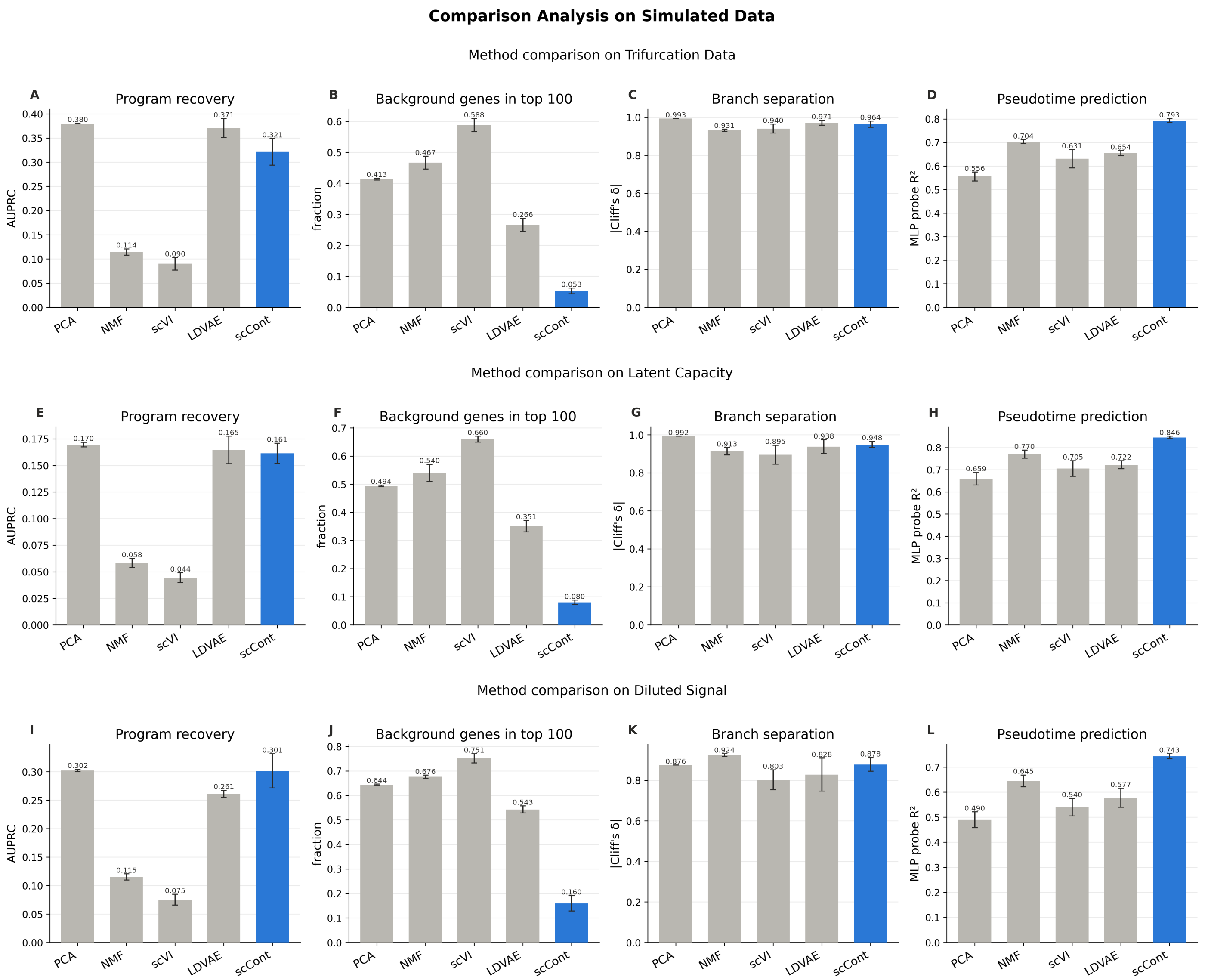


**Figure 2 Comparison of five methods on SERGIO simulated datasets. Panels A-D)** Simulated dataset with trifurcation (3 branches) data. 800 functional genes, 16 functional programs (50 genes per functional program), and 2200 background genes. **Panels E-H) Simulated dataset on Maximum Latent Capacity.** Simulated dataset with latent features at maximum capacity (bifurcation). 800 functional genes, 32 functional programs (25 genes per functional program), and 2200 background genes. **Panels I-L)** Simulated dataset with diluted signal. 400 functional genes, 16 functional programs (25 genes per program), and 2600 background genes. All scCont runs were performed with $k = 10$. Metrics measured are the same metrics in main text Figure 2I-L.

Metabolic Regulation: SLC3A2 (Amino acid transporter), SLC44A2, and GLUL (Glutamine Synthetase) are involved in metabolic reprogramming. The specific enrichment of glutamine metabolism genes suggests a metabolic divergence required to support the alternative cell state found in the bifurcating lineage (20; 34; 24).

### Functional Annotation of Context-Dependent Latent Features

We provide a detailed literature-based annotation for the top-contributing genes identifying the latent features in the A549 datasets under EGF, TGFB1, and TNF stimulation (main text Figure 5).

#### *A549 functional programs*

Cytoskeletal Contractility: ACTB and PDLIM7 are central components of the actin cytoskeleton, facilitating the structural

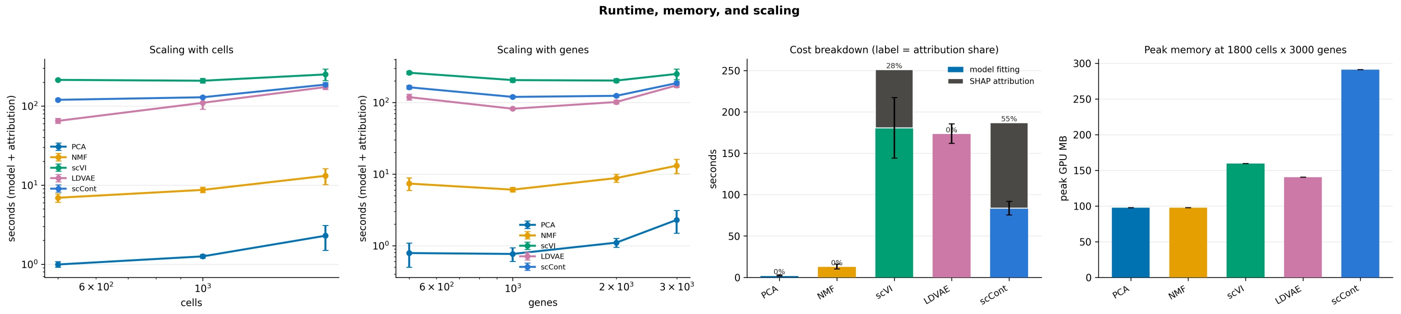


**Figure 3 Runtime analysis on simulated data. A-B)** Mean wall-clock time to produce latent representations and their gene attributions as a function of number of cells at 3000 genes (A) and the number of genes for all cells (B); both axes are logarithmic. **C)** Total runtime cost on the full SERGIO dataset with gene attribution where relevant. Dataset used is the same dataset in main text Figure 2. **D)** Peak memory usage of each model. All models were run five times for each test and the average is reported.

reorganization required for mesenchymal morphology. The co-upregulation of MYL6 (Myosin Light Chain 6) and TPM1 (Tropomyosin 1) specifically indicates the formation of contractile stress fibers, which generate the traction forces necessary for cellular translocation (12; 15; 8).

Keratin Regulation: Similar to the MCF10A model, A549 cells undergo a ”keratin switch” during EMT. However, the specific keratin isoforms upregulated here are distinct from the MCF10A context, highlighting the context-dependent nature of intermediate filament remodeling in response to different inducers (EGF vs. TGFB1) (18).

Microtubule Regulation: Tubulins provide the structural polarity required for directional migration. The upregulation of CCT (Chaperonin Containing TCP-1) genes is critical for the correct folding and assembly of these tubulin subunits, supporting the increased microtubule dynamics observed in the migratory phenotype (5; 3).

Calcium Signaling: The S100 gene family encodes calcium-binding proteins that act as key integrators of signaling and cytoskeletal dynamics. S100 proteins are established prognostic biomarkers in lung cancer, known to promote metastasis by modulating cell stiffness and motility in a calcium-dependent manner (21).

Cell Cycle Proliferation: CCNB1 (Cyclin B1), CCNB2, and CCND1 (Cyclin D1) are critical regulators of cell cycle transitions. Their upregulation in the EGF-induced context suggests a ”Proliferative EMT” or partial state, distinct from the growth arrest typically associated with full TGFB-induced EMT (17; 33; 29).

Integrin Adhesion: Upregulation of Integrin subunits mediates the physical coupling of the cell to the extracellular matrix (ECM). These receptors not only provide traction for migration but also act as outside-in signaling hubs that amplify TGFB1 signaling to sustain the mesenchymal phenotype (30).

Inflammation Signaling (TNF-Specific): CXCL chemokines are potent inflammatory mediators. Their specific upregulation in the TNF-treated condition confirms the activation of NF-$\kappa$B-driven inflammatory pathways, linking the EMT program to an immune-modulatory phenotype often seen in cancer progression (36).

Proteasome Units: The ubiquitin-proteasome system is essential for regulating protein turnover during rapid cellular remodeling. Dysregulation of Proteasome subunits is particularly linked to the regulation of NF-$\kappa$B signaling (via I$\kappa$B degradation), further supporting the inflammatory link in the TNF-induced trajectory (19).

### GO Term Results from Context-Dependency Datasets

As detailed in Supplementary Table 1, the enriched GO terms extracted by scCont reveal distinct regulatory patterns across the context-dependency datasets (2). While the specific enriched genes showed minimal overlap across the twelve distinct conditions, scCont successfully identified that functional pathway overlap is

**Table 1 scCont GO term enrichment across 12 context-dependent EMT datasets.**

| | EGF | TGFB1 | TNF |
|---|---|---|---|
| **A549** | Binding (RNA, cadherin, enzyme), Telomere Maintenance, Protein-folding complexes | Cell adhesion & junctions, Apoptosis, Focal adhesion | Binding (RNA, cadherin), Apoptosis |
| **DU145** | RNA binding, Cell junctions, Focal Adhesion, Metabolic processes, Ribosomal subunits & translation | Binding (protein, RNA), Cytoskeletal components, Cell junctions, Focal Adhesion | Binding (RNA, protein, cadherin), Cell adhesion & junctions, Focal adhesion, Ribosomal components, Mitochondrion/catalytic complexes |
| **MCF7** | RNA binding, Catalytic complex, Ribosomal subunits & translation, Metabolic processes | RNA binding, Apoptosis, Electron transfer activity, Catalytic complex, Ribosomal subunits | Cell adhesion, Catalytic complex, Metabolic processes |
| **OVCA420** | Binding (RNA, protein, cadherin), Cellular junctions, Protein folding | Cellular junctions, Epithelial cell differentiation | Binding (protein, cadherin), Cell adhesion & junctions, Focal adhesion, Ribosomal components and translation |

Note: Datasets from Cook et al. (2) were analyzed to extract enriched spatial clusters (adj. p-val $< 0.05$), corresponding to the methodology shown in main text Figure 1E. GO terms are grouped by their overarching parent term (e.g., cell-cell adhesion is nested under cell adhesion). The complete, unedited list of GO terms is available on the scCont GitHub repository.

predominantly driven by the cell line rather than the EMT-inducing treatment. This aligns perfectly with the main findings of Cook et al. (2), demonstrating that the transcriptomic baseline of the host cell dictates the continuous EMT trajectory more strongly than the external stimulus. For example, scCont consistently detected the enrichment of catalytic complexes and electron transport chain activity across all MCF7 treatments, successfully capturing the established mitochondrial dysfunction characteristic of this specific cell line (13).

### References


1. Naomi E. Butler Tjaden, Stephen R. Shannon, Christopher W. Seidel, Melissa Childers, Kazushi Aoto, Lisa L. Sandell, and Paul A. Trainor. Rdh10-mediated retinoic acid signaling regulates the neural crest cell microenvironment during ens formation. *bioRxiv*, 2025.
2. David P. Cook and Barbara C. Vanderhyden. Context specificity of the EMT transcriptional response. *Nature Communications*, 11(1):2142, 2020.
3. Amanda Cox, Daniel Nierenberg, Oscar Camargo, Eunkyung Lee, Amr S. Khaled, Joseph Mazar, Rebecca J. Boohaker, Tamarah J. Westmoreland, and Annette R. Khaled. Chaperonin containing TCP-1 (CCT/TRiC) is a novel therapeutic and diagnostic target for neuroblastoma. *Frontiers in Oncology*, 12:975088, 2022.
4. Yufang Dai, Jiaqian Zhang, Xiufeng Bai, Shasha Wu, Yanqiong Chen, Dachao Mou, Yun Wang, Yunlong Zhu, and Yi Liu. MFAP5 as a promising biomarker for connective tissue disease-associated interstitial lung disease. *Frontiers in Immunology*, 16:1642408, 2025.
5. Zoltán Dezső, Judith Oestreicher, Amy Weaver, Stephanie Santiago, Sergei Agoulnik, Jesse Chow, Yoshiya Oda, and Yasuhiro Funahashi. Gene expression profiling reveals epithelial mesenchymal transition (EMT) genes can selectively differentiate eribulin sensitive breast cancer cells. *PLoS ONE*, 9(8):e106131, 2014.
6. Jiehui Di, Zhongjun Zhao, Mingyi Xia, Keyu Gao, Keli Chai, Bao Zhu, Wanping Sun, Yanping Zhang, Junnian Zheng, and Yong Liu. Rap2b drives tumorigenesis and progression of colorectal cancer through intestinal cytoskeleton remodeling. *Cell Death & Disease*, 16(1):290, 2025.
7. Amal T. Elhaw, Priscilla W. Tang, Ya-Yun Cheng, Shriya Kamlapurkar, Zaineb Javed, Sarah Al-Saad, Sierra R. White, Ahmed Emam Abdelnaby, Hannah Khan, Alex Seok Choi, Aidan R. Cole, Yeon-Soo Kim, Huda I. Atiya, Mohamed Trebak, Ioannis Zervantonakis, Ronald J. Buckanovich, Katherine M. Aird, Lan G. Coffman, Karthikeyan Mythreye,

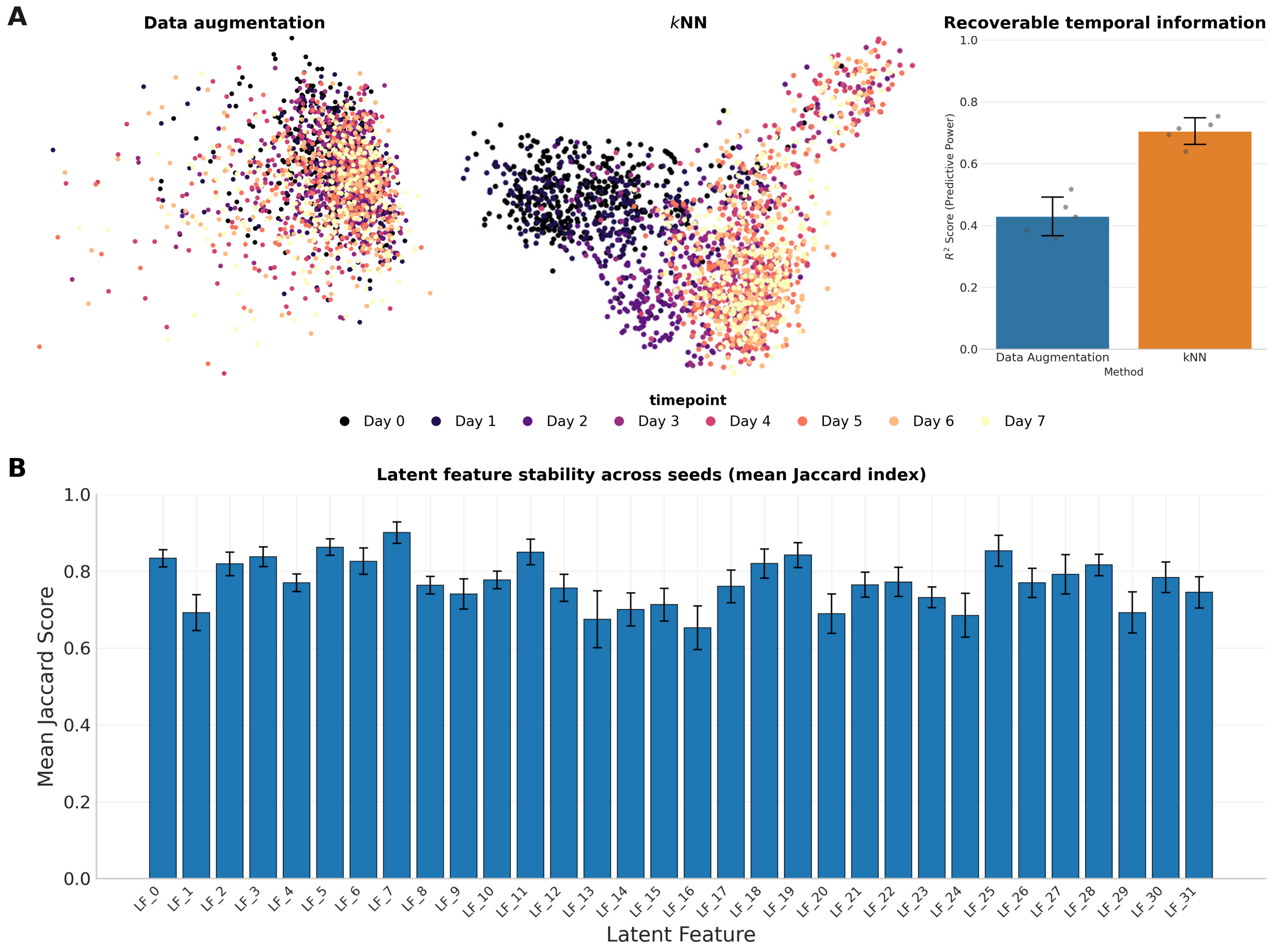


**Figure 4 scCont Algorithm and Parameter Selection. A) k-NN Ablation Study: (Left)** PCA embeddings visualize the latent space generated using standard data augmentation strategies versus the *k*-NN approach. **(Right)** Bar plot displaying the recoverable temporal information ($R^2$ score via 5-fold cross-validation). **B) SHAP Seed Stability:** Robustness of SHAP gene attribution. The vertical bar chart shows the mean Jaccard Index of the resulting gene set for each latent feature across five different initializations. All results were generated using the MCF10A TGFB1 dataset.


and Nadine Hempel. RHOV is a detachment-responsive rho GTPase necessary for ovarian cancer peritoneal metastasis. *bioRxiv*, page 2025.08.12.669944, 2025.

8. Jennifer England, Javier Granados-Riveron, Luis Polo-Parada, Diji Kuriakose, Christopher Moore, J. David Brook, Catrin S. Rutland, Kerry Setchfield, Christopher Gell, Tushar K. Ghosh, Frances Bu'Lock, Christopher Thornborough, Elisabeth Ehler, and Siobhan Loughna. Tropomyosin 1: Multiple roles in the developing heart and in the formation of congenital heart defects. *Journal of Molecular and Cellular Cardiology*, 106:1–13, 2017.

9. Wasundara Fernando, Brianne M. Cruickshank, Raj Pranap Arun, Maya R. MacLean, Hannah F. Cahill, Fiorella Morales-Quintanilla, Cheryl A. Dean, Marie-Claire D. Wasson, Margaret L. Dahn, Krysta M. Coyle, Olivia L. Walker, Melanie R. Power Coombs, and Paola Marcato. ALDH1a3 is the switch that determines the balance of ALDH+ and CD24-CD44+ cancer stem cells, EMT-MET, and glucose metabolism in breast cancer. *Oncogene*, 43(43):3151–3169, 2024.

10. Allison A. Fitzgerald and Louis M. Weiner. The role of fibroblast activation protein in health and malignancy.

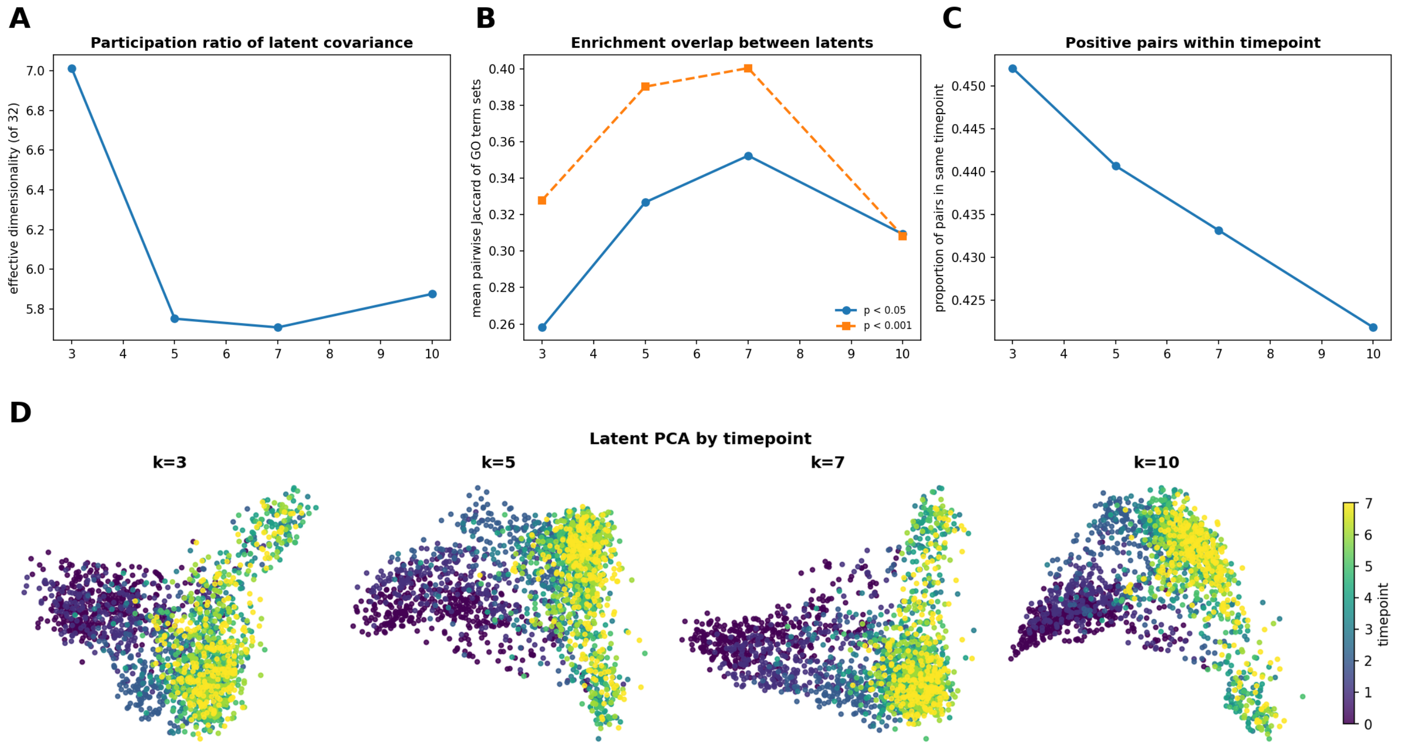


**Figure 5 scCont k sweep on MCF10A TGFB1 dataset. A) Effective dimensionality of the latent space as a function of k.** Participation ratio of the latent covariance. Higher number indicates that latent features are less redundant. **B) Functional overlap between latent features across neighborhood size k.** Plotted value is the mean Jaccard index over all pairs of latent features with at least one enriched term. **C) Temporal composition of positive pairs across neighborhood size k.** Fraction of pairs in which cells are from the same timepoint as a function of $k$. **D) Latent space structure by timepoint for selected k.** Principal component analysis of scCont latent embeddings for $k = 3, 5, 7, 10$. Cells are colored by timepoint.

*Cancer and Metastasis Reviews*, 39(3):783–803, 2020.

11. Jason T George, Mohit Kumar Jolly, Jackson Xu, Jason Somarelli, and Herbert Levine. Survival outcomes in cancer patients predicted by a partial EMT gene expression scoring metric. *Cancer Research*, 77(22):canres.3521.2016, 2017.
12. Yuxi Gu, Shouyi Tang, Zhen Wang, Luyao Cai, Haosen Lian, Yingqiang Shen, and Yu Zhou. A pan-cancer analysis of the prognostic and immunological role of $\beta$-actin (ACTB) in human cancers. *Bioengineered*, 12(1):6166–6185, 2021.
13. M Guha, S Srinivasan, G Ruthel, A K Kashina, R P Carstens, A Mendoza, C Khanna, T Van Winkle, and N G Avadhani. Mitochondrial retrograde signaling induces epithelial–mesenchymal transition and generates breast cancer stem cells. *Oncogene*, 33(45):5238–5250, 2014.
14. Yongsheng Huang, Wenyi Guo, Yuan Zeng, Xinrong Wang, Bohao Fan, Ying Zhang, Lei Yan, Gangli Gu, and Zhao Liu. Identification and validation of a gap junction protein related signature for predicting the prognosis of renal clear cell carcinoma. *Frontiers in Oncology*, 14:1354049, 2024.
15. Xinyuan Jiang, Zhiyong Xu, Sujing Jiang, Huan Wang, Mingshu Xiao, Yueli Shi, and Kai Wang. PDZ and LIM domain-encoding genes: Their role in cancer development. *Cancers*, 15(20):5042, 2023.
16. Briana Lee, Alvaro Villarreal-Ponce, Magid Fallahi, Jeremy Ovadia, Peng Sun, Qian-Chun Yu, Seiji Ito, Satrajit Sinha, Qing Nie, and Xing Dai. Transcriptional mechanisms link epithelial plasticity to adhesion and differentiation of epidermal progenitor cells. *Developmental Cell*, 29(1):47–58, 2014.
17. Bin Li, Jianhua Cheng, Hongyun Wang, Sida Zhao, Haibo Zhu, Chuzhong Li, Yazhuo Zhang, and Peng Zhao. CCNB1 affects cavernous sinus invasion in pituitary adenomas through the epithelial–mesenchymal transition. *Journal of Translational Medicine*, 17(1):336, 2019.

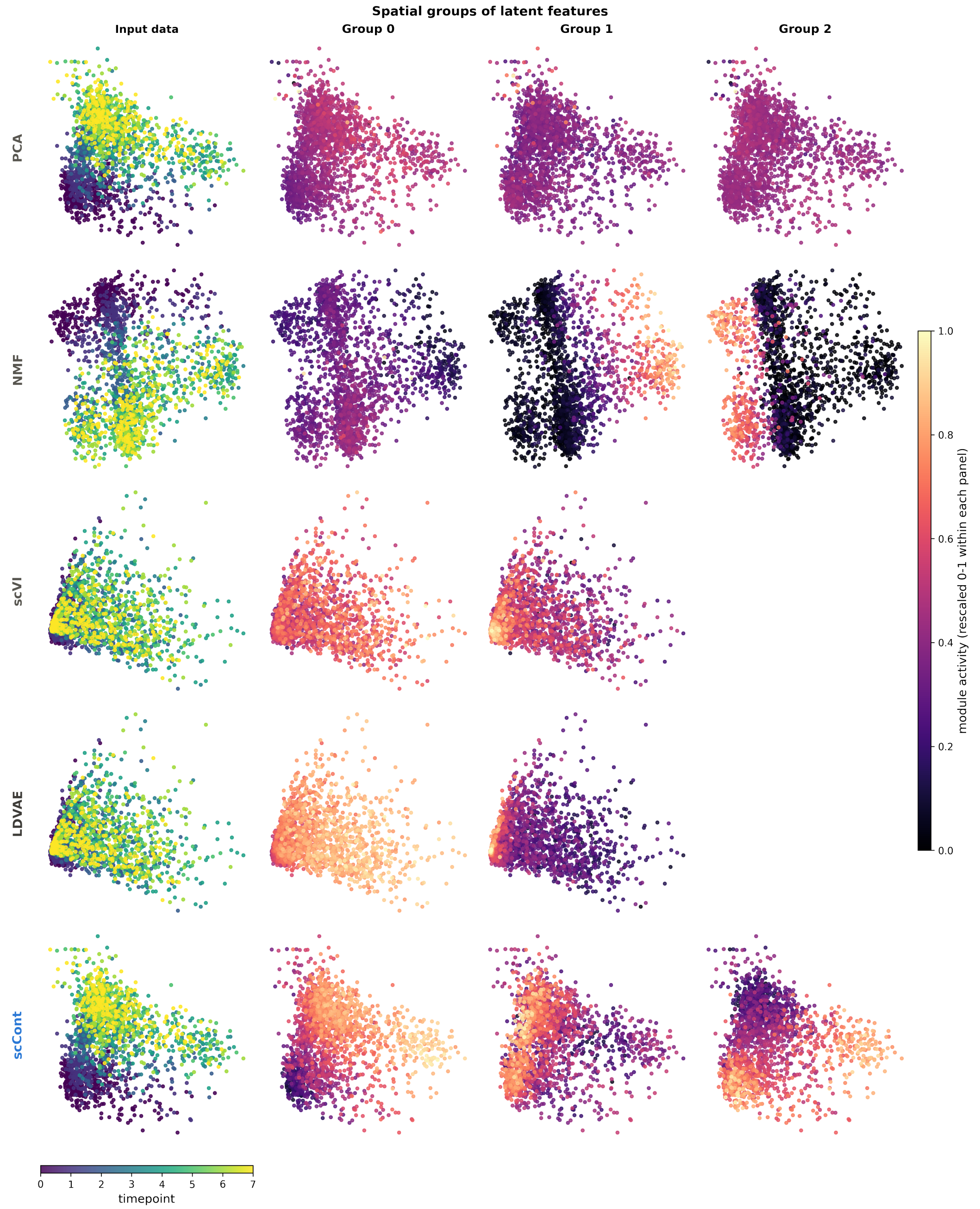


**Figure 6 Qualitative Comparison of Latent Feature Activation.** PCA embeddings of the input data for PCA, NMF, scVI, LDVAE, and scCont ordered from top to bottom. The leftmost column displays the cells colored by experimental timepoint. The subsequent columns visualize the module activity intensity of distinct latent feature cluster groups projected onto the same PCA space. Modules were identified using the method described in main text Figure 1C. To ensure a viable comparison, all models were trained on the MCF10A TGFB1 dataset using the identical set of 3,000 highly variable genes, with variations only in the necessary input normalization steps (see main text Methods for details).

18. Gang Li, Jinbao Guo, Yunfei Mou, Qingsong Luo, Xuehai Wang, Wei Xue, Ting Hou, Tianyang Zeng, and Yi Yang. Keratin gene signature expression drives epithelial-mesenchymal transition through enhanced TGF-$\beta$ signaling pathway activation and correlates with adverse prognosis in lung adenocarcinoma. *Heliyon*, 10(3):e24549, 2024.

19. Na Li and Xianquan Zhan. Integrated genomic analysis of proteasome alterations across 11,057 patients with 33 cancer types: clinically relevant outcomes in framework of 3p medicine. *EPMA Journal*, 12(4):605–627, 2021.

20. Yang Li, Feng Mo, Hua Mu, Jie Zhi, Zhifei Xin, Wujie Zhao, Qingxia Li, and Yitao Jia. Metabolic communication in

colorectal cancer liver metastasis: The role of l-glutamine and slc3a2. *International Immunopharmacology*, 162:115115, 2025.

21. Xiaojie Liang, Xiaoshan Huang, Zihong Cai, Yeling Deng, Dan Liu, Jiayi Hu, Zhihao Jin, Xinyu Zhou, Hongsheng Zhou, and Liang Wang. The s100 family is a prognostic biomarker and correlated with immune cell infiltration in pan-cancer. *Discover Oncology*, 15(1):137, 2024.

22. Paola Marcato, Cheryl A. Dean, Rong-Zong Liu, Krysta M. Coyle, Moamen Bydoun, Melissa Wallace, Derek Clements, Colin Turner, Edward G. Mathenge, Shashi A. Gujar, Carman A. Giacomantonio, John R. Mackey, Roseline Godbout, and Patrick W.K. Lee. Aldehyde dehydrogenase 1a3 influences breast cancer progression via differential retinoic acid signaling. *Molecular Oncology*, 9(1):17–31, 2015.

23. M. Peter Marinkovich. Laminin 332 in squamous-cell carcinoma. *Nature Reviews Cancer*, 7(5):370–380, 2007.

24. Nioosha Nekooie Marnany, Alwyn Dady, Isabelle Duband-Goulet, Frédéric Relaix, Roberto Motterlini, Roberta Foresti, Sylvie Dufour, and Jean-Loup Duband. Non-canonical nuclear function of glutaminase cooperates with wnt signaling to drive EMT during neural crest development. *Nature Communications*, 16(1):5554, 2025.

25. David J. McConkey, Woonyoung Choi, Lauren Marquis, Frances Martin, Michael B. Williams, Jay Shah, Robert Svatek, Aditi Das, Liana Adam, Ashish Kamat, Arlene Siefker-Radtke, and Colin Dinney. Role of epithelial-to-mesenchymal transition (EMT) in drug sensitivity and metastasis in bladder cancer. *Cancer and Metastasis Reviews*, 28(3-4):335–344, 2009.

26. Holly Merta, Kaitlynn Gov, Tadamoto Isogai, Blessy Paul, Achinta Sannigrahi, Arun Radhakrishnan, Gaudenz Danuser, and W. Mike Henne. Spatial proteomics of ER tubules reveals CLMN, an ER-actin tether at focal adhesions that promotes cell migration. *Cell Reports*, 44(4):115502, 2025.

27. Ayalur Raghu Subbalakshmi, Sarthak Sahoo, Prakruthi Manjunatha, Shaurya Goyal, Vignesh A Kasiviswanathan, Yeshwanth Mahesh, Soundharya Ramu, Isabelle McMullen, Jason A. Somarelli, and Mohit Kumar Jolly. The ELF3 transcription factor is associated with an epithelial phenotype and represses epithelial-mesenchymal transition. *Journal of Biological Engineering*, 17(1):17, 2023.

28. Chanjun Sun, Kaishang Zhang, Chen Ni, Jiajia Wan, Xixi Duan, Xiaohan Lou, Xiaohan Yao, Xiangnan Li, Ming Wang, Zhuoyu Gu, Pengyuan Yang, Zhenzhen Li, and Zhihai Qin. Transgelin promotes lung cancer progression via activation of cancer-associated fibroblasts with enhanced IL-6 release. *Oncogenesis*, 12(1):18, 2023.

29. Nicholas P Tobin, Andrew H Sims, Katja L Lundgren, Sophie Lehn, and Göran Landberg. Cyclin d1, id1 and EMT in breast cancer. *BMC Cancer*, 11(1):417, 2011.

30. Donatella Valdembri and Guido Serini. The roles of integrins in cancer. *Faculty Reviews*, 10:45, 2021.

31. Katarzyna Wieczorek, Magdalena Wiktorska, Izabela Sacewicz-Hofman, Joanna Boncela, Andrzej Lewiński, M. Anna Kowalska, and Jolanta Niewiarowska. Filamin a upregulation correlates with snail-induced epithelial to mesenchymal transition (EMT) and cell adhesion but its inhibition increases the migration of colon adenocarcinoma HT29 cells. *Experimental Cell Research*, 359(1):163–170, 2017.

32. Brigham C. Willis and Zea Borok. Tgf-$\beta$-induced emt: mechanisms and implications for fibrotic lung disease. *American Journal of Physiology-Lung Cellular and Molecular Physiology*, 293(3):L525–L534, 2007. PMID: 17631612.

33. Shuai Wu, Rui Su, and Hongyan Jia. Cyclin b2 (CCNB2) stimulates the proliferation of triple-negative breast cancer (TNBC) cells in vitro and in vivo. *Disease Markers*, 2021(1):5511041, 2021.

34. Ying Yang, Longlong Zheng, Jiaxing He, Tao Wu, Haicheng Yang, Bo Zhang, Shuai Zhou, Yueyue Lu, Xianli He,

Jibin Li, and Nan Wang. SLC44a2 negatively regulates mitochondrial fatty acid oxidation to suppress colorectal progression by blocking the MUL1-CPT2 interaction. *Cell Death & Disease*, 16(1):468, 2025.

35. Yingying Zhang, Chan Gao, Yixuan Li, Qinjiao Fu, Yanzhu Liu, Nan Mo, Juanqing Yue, and Ying Wang. Piezo-type mechanosensitive ion channel component 1: a mechano-bioenergetic transducer in the tumour microenvironment. *Annals of Medicine*, 58(1):2603022, 2026.

36. Chenjia Zhou, Ying Gao, Peilun Ding, Tao Wu, and Guang Ji. The role of CXCL family members in different diseases. *Cell Death Discovery*, 9(1):212, 2023.

37. Hongkai Zhuang, Zixuan Zhou, Zuyi Ma, Zhenchong Li, Chunsheng Liu, Shanzhou Huang, Chuanzhao Zhang, and Baohua Hou. Characterization of the prognostic and oncologic values of ITGB superfamily members in pancreatic cancer. *Journal of Cellular and Molecular Medicine*, 24(22):13481–13493, 2020.
